# BORIS/CTCFL-mediated chromatin accessibility alterations promote a pro-invasive transcriptional signature in melanoma cells

**DOI:** 10.1101/2022.02.11.479460

**Authors:** Roy Moscona, Sanne Marlijn Janssen, Mounib Elchebly, Andreas Ioannis Papadakis, Eitan Rubin, Alan Spatz

## Abstract

Melanoma is the deadliest form of skin cancer, due to its tendency to metastasize early. Brother of Regulator of Imprinted Sites (BORIS), also known as CCCTC binding factor-Like (CTCFL), is a transcription regulator that becomes ectopically expressed in melanoma. We recently showed that BORIS contributes to melanoma phenotype switching by altering the gene expression program of proliferative melanoma cells in favor of a more invasive phenotype. However, how BORIS alters the transcriptome remains unclear. Here, ATAC-seq was used to study BORIS-mediated chromatin accessibility alterations in proliferative melanoma cells. Genes that gained promoter accessibility following ectopic BORIS expression, were enriched for melanoma-specific invasive genes as well as invasion-associated biological processes, while promoters of genes associated with proliferation show reduce accessibility. Integration of ATAC-Seq and RNA-Seq data demonstrates that increased chromatin accessibility is associated with transcriptional upregulation of genes involved in tumor progression processes, and the aberrant activation of oncogenic transcription factors, while reduced chromatin accessibility and downregulated genes, were associated with repressed activity of tumor suppressors. Together, these findings indicate that BORIS mediates transcriptional reprogramming in melanoma cells by altering chromatin accessibility and gene expression, shifting the cellular transcription landscape of proliferative melanoma cells towards a pro-invasive genetic signature.

**Significance:** We recently reported that BORIS contributes to melanoma phenotype switching by altering the transcriptional landscape of melanoma cells from a proliferative to an invasive state. In this study, using ATAC-Seq, we demonstrate that ectopic BORIS expression in proliferative melanoma cells leads to increased chromatin accessibility at promoters of upregulated invasion-associated genes. Importantly, by integrating the ATAC-Seq data with RNA-Seq data, we were able to identify key cancer-associated transcription factors that become aberrantly activated or repressed following ectopic BORIS expression. Taken together, this study sheds light on the mechanisms by which BORIS mediates phenotype switching in melanoma cells.

## 1 INTRODUCTION

Primary cutaneous melanoma accounts for most of the deaths from skin cancer, due to its ability to quickly disseminate and form distant metastases. Worldwide, approximately 230,000 new cases of melanoma are diagnosed each year, resulting in about 55,000 deaths (Siegel et al. 2020). In the progression from primary tumor towards an aggressive metastatic stage, cells undergo a reversible transition between a proliferative and invasive state. This transition is known as “phenotype switching”, which resembles epithelial-mesenchymal transition (EMT) as observed in cancer cells with epithelial origin (Hoek et al. 2008a; Hoek and Goding 2010). Studies of melanoma identified specific genes and transcription factors (TFs) involvement in different stages of phenotype switching (Hoek et al. 2006; Widmer et al. 2012; Jeffs et al. 2009; Tirosh et al. 2016). The ‘proliferative’ state of melanoma cells is linked to high expression levels of TFs such as the melanocyte differentiation gene *MITF* (Hoek et al. 2008b), *SOX10* (Shakhova et al. 2012) and *PAX3* (He et al. 2005). On the contrary, melanocytes of the ‘invasive’ signature express low levels of ‘proliferative’ signature TFs and higher levels of invasion-related TFs such as *TEAD* (Verfaillie et al. 2015), *BRN2* (Pinner et al. 2009), Activator Protein-1 (AP-1) family (Jochum et al. 2001), *NFATC2* (Perotti et al. 2019) and *NFIB* (Fane et al. 2017b).

Epigenetic studies of melanoma genesis revealed chromatin landscape changes in different phenotypic states of the disease (Dahl and Guldberg 2007; Fiziev et al. 2017; Verfaillie et al. 2015). The genetic and epigenetic alterations are driven by transcriptional regulators. Exploring their exact mechanism of action in melanoma is pivotal for our understanding of the development of the disease. An important chromatin modifier in malignancies is Brother of Regulator of Imprinted Sites (*BORIS*), also known as CCCTC binding factor-Like (*CTCFL*). It influences the chromatin landscape and transcription, both directly and indirectly. Despite the study of *BORIS’s* role in various cancer types, its exact mechanism of action in melanoma development and progression is yet to be deciphered.

BORIS is a DNA-binding protein with high similarity to its ‘sibling’ gene *CTCF* (Loukinov et al. 2002), the multifunctional TF, known as the ‘master weaver’ of the genome (Phillips and Corces 2009). Under normal conditions, BORIS protein functional expression is strictly restricted to the testis, whilst it becomes aberrantly expressed in different types of cancers (Loukinov et al. 2002), being the most frequently switched among cancer testis antigens (CTAs) (Wang et al. 2016). The delicate balance between BORIS and CTCF expression in the testis is shifted towards *BORIS* in malignant cells (Klenova et al. 2002), while CTCF demonstrates tumor suppressive properties (Kemp et al. 2014; Rasko et al. 2001; Fiorentino and Giordano 2012), BORIS has been reported to act as an oncogene in different malignancies, such as breast, prostate and colorectal carcinoma (Gaykalova et al. 2012; Liu et al. 2017; Renaud et al. 2011; Smith et al. 2009; Dougherty et al. 2008; D’Arcy et al. 2008; Cheema et al. 2014; Zhang et al. 2017). Of note, in ovarian cancer, BORIS promotes invasiveness (Hillman et al. 2019). Similarly, we show in a previous study that BORIS ectopic expression in melanoma cells reduces proliferation and increases invasiveness properties, through a transcriptional and phenotypic switch (Janssen et al. 2020). However, how BORIS mediates these transcriptional changes in melanoma cells remains elusive.

Here, the impact of BORIS ectopic expression on the transcriptional landscape in melanoma cells was explored by assessing chromatin accessibility using the Assay for Transposase-Accessible Chromatin followed by sequencing (ATAC-Seq) (Buenrostro et al. 2013) to reveal genome wide chromatin accessibility changes induced by ectopic BORIS expression. Based on chromatin analysis of the ATAC-Seq data, combined with gene expression data from RNA-Seq, BORIS unique target genes and TFs were revealed. Among the genes identified as BORIS up-regulators, an over-representation of tumor invasiveness and progression factors were identified. In addition, enrichment analysis of genes with a differential accessible promoter region, show high enrichment in invasiveness-related biological processes. Overall, our data indicates a unique BORIS genetic signature in melanoma cells, defined by the genes and TFs it governs in its role as a regulator of chromatin accessibility and gene transcription, in a manner which has the potential to drive tumor progression capabilities of melanoma cells towards metastasis dissemination.

## 2 MATERIALS AND METHOD

### 2.1 Cell lines

Early passage MM057 melanoma cells (from Dr. G. Ghanem, Institut Jules Bordet, Brussels, Belgium) were cultured in Ham’sF10 medium supplemented with 8% heat-inactivated fetal bovine serum, 1% penicillin-streptomycin and 1% GlutaPlus. Doxycycline (dox) inducible cells were selected by the addition of 150 μg/ml G418 (Neomycin) for a minimum of 2 weeks or until non-infected control cells were killed. Upon selection, MM057 cells were maintained in the presence of 50 μg/ml G418. Doxycycline (dox; Clontech) was added to the medium for the indicated dose and time and refreshed every 48 h. Bimonthly tests for mycoplasma demonstrated the absence of contamination. All cultures were maintained at 37 °C in a 5% CO2 humidified atmosphere.

### 2.2 Generation of expression vectors

Full-length human BORIS cDNA was amplified by PCR on RNA extracted from HEK293T cells using iProof (Bio-Rad) with primers containing BamHI and MluI restriction sites (forward 5′-CAGCGGATCCACTGAGATCTCTGTCCTTTCTGAG-3′ and reverse 5′-GCGGACGCGTCT CACTTATCCATCGTGTTGAGGAGCATTTCACAGG-3′). The PCR product was digested with BamHI and MluI (Fermentas FastDigest) and inserted into pDONR221 (Invitrogen cat # 12536017) by a BP reaction (GatewayTM BP ClonaseTM II Enzymemix, Invitrogen) to generate pENTR-BORIS with six C-terminal His-tags followed by a stop codon. To generate pI20-BORIS-6xH the entry clone was recombined into the lentiviral destination vector pI20 (empty vector (EV)-6xH, a kind gift from Stephen Elledge (Meerbrey et al. 2011), Addgene plasmid #44012) using Gateway LR ClonaseTM II (Fisher).

### 2.3 Transfection, lentivirus production, and infection

For virus production, pI20-BORIS-6xH or pI20-EV-6xH lentiviral expression vector (10μg DNA) was transfected into HEK293T cells along with the viral packaging plasmids pSPAX2 (7.5μg DNA) and pMD2.G (5μg DNA) using polyethyleneimine (PEI; Polysciences, DNA:PEI ratio of 1:2.5) and Opti-MEM transfection medium (Invitrogen). The medium was replaced by fresh medium 3h after transfection. At 48h post transfection, virus-containing supernatant was harvested, centrifuged, passed through a 0.45μm syringe filter, and either used directly for infection or stored at -80°C. Cells were infected twice by incubation with a mixture of viral supernatant, medium (1:1 ratio) and polybrene (8 μg/ml, Sigma–Aldrich) for 12h. The viral mixture was replaced by fresh medium for 24h before selection. Expression of the constructs was verified by a combination of immunoblot and quantitative PCR.

### 2.4 Assay for transposase accessible chromatin followed by sequencing (ATAC-Seq)

For ATAC-Seq library preparation cells were detached from the culture dish with citric saline (270mM KCl, 30mM sodium citrate), passed through a 40um filter and counted using a hemocytometer to collect 50.000 cells for all four experimental conditions. To isolate nuclei, cells were first lysed for 20 min on ice in lysis buffer 1 (0.1% Sodium citrate tribasic dehydrate (Sigma, S1804) and 0.1% Triton X-100 (Bioshop, TRX777.100)) followed by 20 min lysis on ice in lysis buffer 2 (10mM Tris-HCl pH7.4, 10mM NaCl, 3mM MgCl, 0.1% Igepal CA630 (Bioshop, NON999.500) and 0.1% Tween-20 (Bioshop, TWN510.500). The nuclei were pelleted by centrifugation for 10min at 600g (4°C) and supernatant was carefully removed to reduce mitochondrial DNA (mtDNA) content. For the transposase reaction, the nuclei pellet was re-suspended in 25μl transposase reaction mix containing of 4μl Tn5 transposase in 1x transposase reaction buffer (Nextera DNA library kit, Illumina, FC-121–1030) and 0.1% Tween-20 (Bioshop, TWN510.500), and incubated for 30min at 37°C with gentle shaking (400rpm). Immediately following transposition, DNA was purified using the MinElute PCR purification Kit (Qiagen, 28004), according to manufacturer’s instructions. Library fragments were amplified in two steps. First, 5 cycles of PCR were performed using NEBNext® High-Fidelity PCR master mix (New England BioLabs, M0541) with 1.25μM barcoded Nextera PCR primers (*Supp* 10.). in a final volume of 50μl on a BioRad C1000 Thermal Cycler. The cycling program used was as follows: 5 min at 72°C, 30 sec at 98°C, 5 times cycling for 10sec at 98°C, 30sec at 63°C and 1 min at 72°C. To reduce GC and size bias during library amplification, 5μl from the PCR mix was used for a 15μl qPCR side-reaction containing the same reagents with the addition of SYBRgreen (9X, Life Technologies, S7563) to determine the number of additional PCR cycles and stop amplification prior to saturation. The CFX96 TouchTM (Bio-Rad, 1855196) qPCR machine was used with the following cycling conditions: 30 sec at 98°C, 20 times cycling for 10sec at 98°C, 30sec at 63°C and 1 min at 72°C followed by melt curve analysis. Using the CFX ManagerTM Software (Bio-Rad), the fluorescent intensity was plotted versus cycle number and the additional number of cycles was established as the cycle number corresponding to 1/4^th^ of the maximum fluorescent intensity. Next, the initial PCR reaction mix was amplified for 3 additional PCR cycles. Library preparation for each experimental condition was performed in triplicate and samples were combined post PCR amplification for quality controls and sequencing. One third of the library was cleaned up using KAPA Pure Beads (1.8X; Roche, 7983298001) and analyzed on a 2.5% agarose (Bio-Rad, 1613101) gel to verify nucleosomal patterning. The remainder of the library was size selected using KAPA Pure Beads (0.55X-1.8X; Roche, 7983298001) to enrich for 150-750 bp fragments. Part of the DNA from the size selected library was diluted 1 in 5 and used for qPCR to confirm enrichment for open chromatin and estimate mtDNA content. The qPCR reaction contained 2.5μl DNA template, 5ul PerfeCTa SYBR Green FastMix (Quanta, 95072-012) and 300nM forward and reverse primer. The thermal cycling program used was as follows: 3 minutes at 95°C, 40 times cycling for 15 seconds at 95°C and 30 seconds at 60°C followed by melt curve analysis. To assess enrichment of open chromatin primers were designed in genomic regions that are often marked by open chromatin (GAPDH and KAT6B) or closed chromatin (SLC22A3 and RHO). A minimum difference in quantitation cycle (Cq) value of 3, representing an 8-fold enrichment, was considered as enrichment of open chromatin in the libraries. In addition, we performed qPCR to approximate the level of mtDNA in our libraries (established from purified and washed nuclei) as well as libraries prepared from whole cell lysate using 2 sets of primers that are referred to as mtDNA and ND1-mtDNA. The sequences of these primers as well as the primers directed at mtDNA are listed in *Supp* 10.). The remaining DNA of the libraries was submitted to the McGill University and Genome Québec Innovation Centre and fragment distribution was assessed by TapeStation (Agilent), libraries were quantified using Quant-iTTM PicoGreen® dsDNA Reagent® (ThermoFisher Scientific), pooled and sequenced paired-end (100bp read length) on one lane of an Illumina HiSeq4000.

### 2.5 A bioinformatics pipeline for extracting open chromatin regions from ATAC-Seq data

To gain insight into the impact of BORIS on chromatin arrangement in melanoma, ATAC-Seq was utilized with two biological replicates over the four experimental conditions: BORpos, BORneg, EVpos, EVneg. ATAC-Sequencing results, totaling in as 8 raw FastQ reads files, were first subjected to QA with FastQC (Andrews et al. 2010). The FastQC output reports (data not shown) indicated several problematic sequencing results in the following fields: (1) per-tile sequence quality and (2) adapter content contamination. Each abnormal finding was addressed before any further analysis. In all sequencing samples, specific tiles with low Q-scores, were identified concentrated around the last 20 bp positions of the 120 bp input DNA-library. Tiles which contained low-quality reads were removed, using the filterbytile.sh script from BBMap tool (Bushnell B. 2014; Bushnell et al. 2017), using default parameters. In addition, an adapter contamination of the Nextera Transposase Sequence was detected across all samples and was removed using the trim_adapters script by atactk: a toolkit for ATAC-Seq data (Parker 2015); default parameters were used. Furthermore, the first 17 bp of all reads were trimmed with cutadapt (Martin 2011) to ensure high base quality. The final reads had a sequence length of 83 bp and an average of 164,561,210 (Range; 164,424,288-187,302,698) paired-end reads were obtained for all four samples conditions.

The trimmed reads were then aligned to the human reference genome (version GRCh38.p12) using BWA-MEM (Li 2013), with the following parameters: *bwa mem -M -a -t 30 hg38*.*fa. read1*.*fastq read2*.*fastq*. The aligned reads were saved as BAM files and further indexed and name-sorted with SAMtools (Li et al. 2009). Next, duplicate alignments were marked and removed with Picard-Tools(Wysoker et al. 2009) MarkDuplicates tool, using the following command; *picard*.*jar MarkDuplicates I=file_name*.*bam O=file_name*.*no_duplicates*.*bam M=file_name*.*txt REMOVE_DUPLICATES=true VALIDATION_STRINGENCY=LENIENT*. The average duplication rate (including PCR and optical duplicates), as determined by MarkDuplicates, was 31.88% (Range; 30.2%-35.1%). After duplication removal, an additional filtering step was implemented with SAMtools, using the following command: *samtools view -b -h -f 2 -f 3 -F 4 -F 8 -F 256 -F 1024 -F 2048 -q 20 -e chrM -e chrY -e ‘VN:’ file_name*.*bam*. At the end of this sifting step, reads mapping to mitochondrial DNA (mtDNA) and Chromosome Y were excluded, in addition to reads that were not mapped/paired properly or mapped to multiple places in the genome. Lastly, reads with Q-score<20 were removed as well. The final autosomal properly aligned sequences, are described in the results section, for downstream analyses were saved in BAM format. The correlation between each of the two replicates of each one of the four conditions (Figure S1) was calculated with deepTools2 (Ramírez et al. 2016) using the following two commands:*(1)multiBamSummary bins --bamfiles BAM_file_1*.*bam BAM_file_2*.*bam -o Condition*.*npz -p “max” --ignoreDuplicates --minMappingQuality 20* ; *(2) plotCorrelation --corData Condition*.*npz --corMethod spearman --whatToPlot scatterplot --labels Condition _1 Condition _2 -o Condition _name*.*pdf --outFileCorMatrix Condition_file_name*. The sifted BAM served as input for MACS2 (Feng et al. 2012; Zhang et al. 2008).

Genomic regions with an abundance of transposition events detected with MACS2 were considered open chromatin regions (OCRs). The MACS2 call-peak function was used with the following command, customized for ATAC-Seq reads: *macs2 callpeak -g hs -q 0*.*05 -f BAMPE -B*. The results of MACS2 were saved as BedGraph and narrowPeak files, that were further used to find enrichment in ATAC-Seq signal based on pooled, normalized data. In addition, MACS2 output provides Excel files that contain all peaks location, length, enrichment and statistic-index values. Blacklisted regions as defined by the ENCODE file ENCFF419RSJ.bed, were removed.

### 2.6 ATAC-seq data analysis of differential chromatin accessibility regions with R programing

The sifted reads (as BAM files and the MACS2 output narrowPeak files) served as input for the analyses in a dedicated script developed for this purpose in the R Programming language (R Core Team 2015) with Bioconductor (Huber et al. 2015) packages. First, the distribution of the final reads was examined across all chromosomes. In a global view, ATAC-Seq reads were distributed proportionally to the chromosomes size. ATAC-Seq should represent a mix of fragment lengths corresponding to nucleosome-free positions (NFRs) (fragments length<100bp), mono-nucleosome (fragments length: 180-247bp) and poly-nucleosome fractions (fragments length>315bp), as defined by the Greenleaf research team (Buenrostro et al. 2013; Schep et al. 2015). By plotting the insert size (Figure S6), the typical ATAC-Seq distribution was verified, results displayed the expected distribution pattern. All samples were of the desired insert size distribution, with the NFRs representing the bulk of reads, while the number of reads decreased as insert size increased. Next, ATAC-Seq fragment reads >100bp were discarded, since they do not represent NFRs. NFR fragments reads of length <100bp were then plotted in a PCA (Figure S2). Non-redundant peaks were defined as peaks present in both replicates (two samples) of the same tested condition. Only peaks which met this criterion were selected for further assessment of differential alterations in the ATAC-seq signal. The sifted ATAC-Seq reads were saved in narrowPeak format and used for downstream analyses. This file contains information about the ATAC-Seq peaks, such as start and end of each peak as well as the value for the enrichment signal and statistical index. Peaks were annotated to their nearest corresponding gene. In addition, a genomic region was assigned to each peak. The package ChIPseeker (Yu et al. 2015) in R software was used for the genomic assignments and annotations. Differential accessible regions (DARs) analysis was performed, to evaluate differences in ATAC-seq signals between EVpos and BORpos samples. Peaks which mapped to promoters (0-3kb from annotated transcription start site (TSS) as defined in the hg38 reference genome) and found to be differentially accessible between the two tested conditions were defined as DARs (Figure 1B). Ambiguous 196 DARs which showed significantly increased and decreased accessibility of the promoter region of genes to which they were mapped, were excluded from any further analyses. To define accessible chromatin regions as statistically significant, a false discovery rate (FDR) cutoff (FDR<0.05) was used. Accessible chromatin regions that met the cutoff were defined as DARs. Consequentially, this directly influenced the definition of genes the DARs were annotated to, as this cutoff defined the significant genes with a DAR mapped to their promoter.

**Figure 1.**
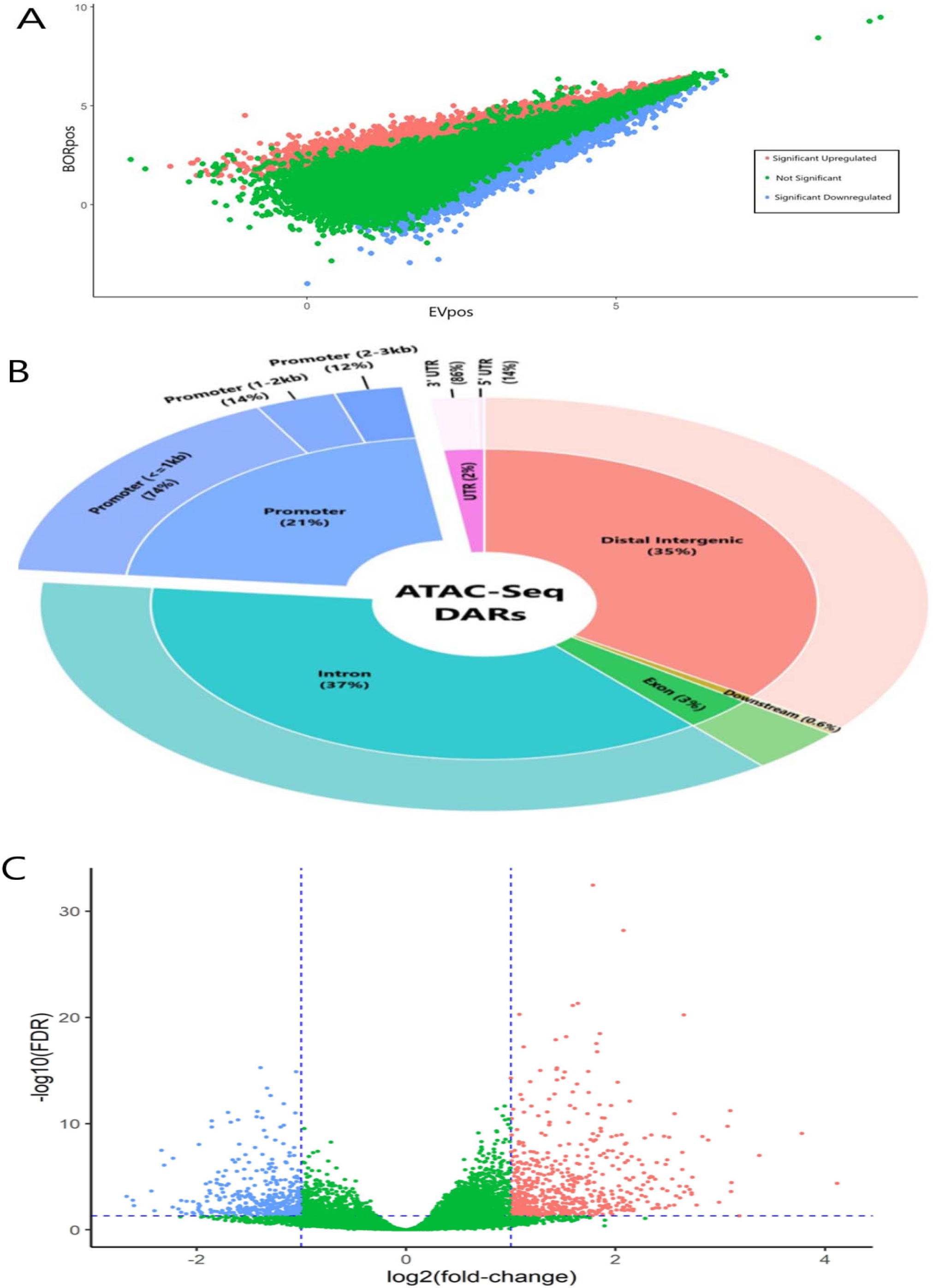
Ectopic BORIS expression in melanoma cells leads to significant differential alterations in chromatin accessibility, especially at promoters. **(A-C)** BORIS expression was induced in the MM057 melanoma cells. Chromatin from cells nuclei was sequenced with ATAC-Seq. Results of a differential analysis of ATAC-seq peaks of Nucleosome Free Regions (NFRs) within Open Chromatin Regions (OCRs), of EVpos vs BORpos samples are shown. **(A)** Fragments Per-Kilobase per-Million mapped fragments (FPKM) values of all identified peaks, for the Evpos and BORpos tested condition. Each dot in the scatter plot represents one out of the 158,534 peaks identified and its value is the average FPKM value of two biological replicates of the two tested condition. FPKM values were calculated from ATAC-Seq BAM files with DESeq2 function ‘fpkm’ in R programing, for each repeat of a tested condition separately: BORpos_1, BORpos_2, EVpos_1, EVpos_2. The values were log_2_ transformed and the two replicates of the same condition were averaged, since they represent repeats of the same tested condition on the same cell line and are highly correlated. Non-significant peaks which did not meet the defined threshold (FDR<0.05 and |log_2_FC|>0) are colored in green, while the 22,594 significant peaks are colored in red (12,010 upregulated {log_2_FC>0}). Blue peaks (10,595 downregulated {log2FC<0}) are those defined as Differentially Accessible Regions (DARs). **(B)** Genomic distribution of DARs. DARs were defined as statistically significant (FDR<0.05) ATAC-seq peaks of NFRs within OCRs. DARs were annotated to the closest genomic region, within a window around a gene’s Transcription Start Site (TSS). The pie chart depicts the proportion of 22,594 DARs, across the human genome (GRCh38), and the assigned genomic regions of each DAR. The inner circle parentheses next to each genomic region, are the percentage of the region out of all DARs. The outer circle parentheses of promoter and UTR regions, represent the percentage of each sub-region out of the whole parent region term. The outer circle parentheses next to promoter regions represent the distance from TSS. R programing language with Bioconductor package ChIPseeker with annotatePeak function was used to characterize the annotation distribution of peaks to genomic regions. **(C)** Volcano plot of the proportion of ATAC-Seq peaks annotated to promoters (0-3kb from TSS). All peaks annotated to promoters were further annotated to the promoter’s corresponding gene. In total, all promoter peaks were annotated to 16,161 genes; each dot in the volcano plot represents one gene. Colored in green are genes which did not meet two cutoffs: FDR<0.05 and|log_2_FC|>1, while colored in red are 651 upregulated (log_2_FC>1) and blue 393 downregulated (log_2_FC<-1) statistically significant (FDR<0.05) genes which had a promoter region with increased (red) or decreased (blue) chromatin accessibility and defined as DAR. Blue dashed lines; X-axis at X=1, X=-1. Y-axis, Y=1.30102999566 (FDR=0.05). For the differential analysis, Bioconductor DESeq2 was used. ATAC-Seq, Assay for Transposase Accessible Chromatin followed by Sequencing; BORpos, BORIS-vector samples positive to doxycycline; Evpos, empty-vector samples positive to doxycycline; FC, fold-change; FDR, false discovery rate.

### 2.7 Functional analysis of significant promoter DARs and differentially expressed genes with Genomic Regions Enrichment of Annotations Tool (GREAT)

Here, promoter DARs were defined as significant (FDR<0.05) OCRs ATAC-Seq peaks mapped to promoters (0-3kb from annotated TSS). The aim of this analysis was to functionally characterize the biological meaning of genomic regions defined as promoter DARs. The DARs list was divided into those with increased (upregulated) versus decreased (downregulated) promoter accessibility. The upregulated promoters consisted of 3,573 genomic region peaks with log_2_FC>0, while the downregulated promoters included 1,246 genomic region peaks with log_2_FC<0. The gene-list of upregulated DARs & differentially expressed genes (DEGs) genomic regions consisted of 554 genomic region peaks with log_2_FC>0. All lists of genomic regions were saved as BED files and served as input to find enriched gene ontology biological processes (GO-BP) terms among promoters DARs, using Genomic Regions Enrichment of Annotations Tool in R (rGREAT software, version 4) (Zuguang Gu 2020; McLean et al. 2010). The rGREAT analysis aims to associate biological functions to genomic regions. For statistical analysis, the rGREAT output of hypergeometric test results were considered, as they provide gene coverage and FDR values, for each tested GO-BP term.

### 2.8 Gene Set Enrichment Analysis with (GSEA) using the ATAC-Seq based gene-list

The GSEA tool (Subramanian et al. 2005) was used. For this analysis, all identified genes with an ATAC-Seq peak mapped to their promoter (0-3kb from annotated TSS) region, without any statistical cutoff, were used, as recommended by GSEA. This analysis aimed to directly compare between ATAC-Seq identified genes to RNA-Seq-identified gene list, established previously by us (Janssen et al. 2020). The ATAC-Seq gene list (16,160 genes) was pre-ranked by each gene’s promoter accessibility log_2_FC value. The generated gene list served as input for the GSEA-pre-ranked tool, with 10^4^ permutations, using the exact same analysis protocol as used with the RNA-Seq data. As the direct comparison between the ATAC-Seq-based GSEA results to the RNA-Seq based GSEA results was required, the same gene sets were tested, i.e., the GO-BP gene set and hallmark EMT gene sets from the Molecular Signatures Database (Liberzon et al. 2015), additionally, the melanoma invasiveness gene set of (Verfaillie et al. 2015) was used. FDR<0.05 cutoff defined statistically significant enrichment among the tested gene-sets.

### 2.9 Differential transcription factor activity analysis with diffTF

A computational framework called diffTF (Berest et al. 2019) was used to identify differential TFs activity based on chromatin accessibility data (ATAC-Seq). This tool also uses as input genome-wide predicted TFs binding sites sequences in the form of position weight matrix (PWM). The addition of a third layer, in the form of gene expression (RNA-seq), enables diffTF to classify TFs as genetic activators or repressors, if the expression level of the TFs target genes, has changed concordantly with chromatin accessibility around the TFs binding sites. For PWM data the PWMscan (Ambrosini et al. 2018) was used with the defined database of HOCOMOCO (Kulakovskiy et al. 2018) (version 11), which houses a collection of 769 TF binding motifs. DiffTF default settings were used, the signal of ATAC-Seq reads around each TF binding sites was extended by 100 bp in each direction and a fold-change of chromatin accessibility was calculated, based on a comparison between EVpos and BORpos. Bootstrapping of 10^3^ was used as recommended, to account for differences in GC content at the different binding sites. Weighted mean difference is the index for ΔTF activity, where a positive value indicates increased TF activity (either as an activator or a repressor), while negative values indicate a less active TF in the BORpos sample as compared to EVpos. Here, only TFs with positive weighted mean difference values were investigated since they indicate TF with increased activity (either as an activator or a repressor) in response to ectopic BORIS expression. To validate that the detected differential TFs activities arise due to ectopic BORIS expression, an additional comparison was made between EVneg and EVpos samples. This comparison did not yield any overlapping results in TFs activity directionality (activators or repressors) with the main comparison of EVpos and BORpos. Indicating that the observed transcriptional alterations were occurring in BORpos cells only, as a result of ectopic BORIS expression.

## 3 RESULTS

### 3.1 Identification of differentially accessible regions in melanoma cells expressing ectopic BORIS

To study the effect of ectopic BORIS expression on chromatin accessibility, ATAC-Seq was performed in duplicate on samples of MM057 melanoma cell line bearing a BORIS-encoding (BOR) or Empty (EV) Dox-inducible vector, grown in the presence (pos) or absence (neg) of Dox. ATAC-Seq results for all samples displayed high mappability to the human genome with 81.77%±8.3 (mean±standard deviation (sd)) mapped reads, which translated to 48,758,922±22.9*10^6^ final properly paired reads for downstream analyses. All ATAC-Seq replicates showed a high correlation for the number of reads mapping to the same location in the human genome (R2=0.98±4.10^−4^) (Figure S1), demonstrating that ATAC-Seq results are highly reproducible. Furthermore, a Principal Component Analysis (PCA) representing the 8 ATAC-Seq biological repeats separated samples according to BORIS expression, with BORpos samples clustering together, apart from the control samples, on PC1 accounting for 85% of the observed variance between samples. PC2 accounts for a minor 7% of observed variance and samples on it cluster by a small batch effect (Figure S2). These findings confirm that the vast majority of variance between samples is due to BORIS over-expression and not to a response to Dox or empty vectors, or culture conditions.

To identify DARs between EVpos and BORpos, ATAC-Seq peaks of OCRs from NFRs for each experimental condition were determined. Next, a differential accessibility analysis was performed, aimed at identifying changes in chromatin accessibility of OCRs peaks between EVpos and BORpos. This initial analysis identified 158,534 differential peaks between the two tested conditions. Correction for multiple testing (FDR<0.05), classified 22,594 of the peaks as significant and those were defined as DARs (Figure 1A). Annotation of the DARs to genomic regions revealed that most DARs were in introns (37%), distal intergenic regions (35%) or promoter regions (21%) (Figure 1B). This distribution did not differ from the distribution of all identified 158,534 peaks (data not shown). Annotation of all identified peaks to their corresponding gene, resulted in assignment to a total of 20,255 known genes. Of these, 9,427 were assigned to peaks that were defined as statistically significant (FDR<0.05) (i.e., DARs) in the differential analysis. To confirm that the DARs identified here are specific to ectopic BORIS expression and not due to the presence of Dox, ATAC-Seq data of EVneg were compared to those of EVpos. No statistically significant chromatin regions were identified between EVneg and Evpos (Figure S3), indicating that the reported DARs are indeed due to ectopic BORIS expression.

### 3.2 Ectopic BORIS expression leads to increased chromatin accessibility at promoter regions

To determine whether ectopic BORIS expression preferentially leads to a gain or loss in chromatin accessibility in MM057 melanoma cells, the fold change in chromatin accessibility between EVpos and BORpos cells was analyzed. Out of all 22,594 DARs mapped to any genomic location, 5,293 DARs exhibited increased chromatin accessibility (log_2_FC>1), while 5,347 DARs showed decreased chromatin accessibility (log_2_FC<-1). Given the role of BORIS in transcriptional regulation at promoters (Bergmaier et al. 2018), we focused on DARs which were mapped to the promoter region of genes (0-3 kb from the annotated TSS). This analysis revealed 4,812 statistically significant (FDR<0.05) DARs in promoter regions. Interestingly, 16% of these promoter DARs showed an increase in chromatin accessibility (log_2_FC>1, N=792), while 11% had reduced chromatin accessibility (log_2_FC<-1, N=543) (Figure 1C). These findings suggest that ectopic BORIS expression preferentially contributes to increased chromatin accessibility at promoters, as compared to all other genomic regions. BORIS’ own promoter ranked at the top of the list of promoters DARs, with a log_2_FC of 5.47, which translates to a 44-fold increment in accessibility of the *BORIS* promoter in BORpos compared to EVpos control cells.

To gain insight into potential functional consequences of altered chromatin accessibility at promoter regions, we tested enrichment of gene-sets present in the GO-BP database among DARs mapped to promoters (0-3kb), using the rGREAT software (Zuguang Gu 2020; McLean et al. 2010). This analysis revealed 57 biological processes as statistically significant (FDR<0.05) among 3,570 DARs mapped to promoters with increased chromatin accessibility (log_2_FC>0), while 89 biological processes were identified as statistically significant (FDR<0.05) among 1,242 DARs mapped to promoters with reduced chromatin accessibility (log_2_FC<0). Among the positively enriched biological processes, some tumor progression processes such as ‘substance dependent cell migration, cell extension’ and ‘regulation of stem cell differentiation’, are present (Figure 2), corroborating with the suggested role of BORIS in cancer stemness (Garikapati et al. 2017; Alberti et al. 2015). Prominent negatively enriched processes include ‘epithelial cell proliferation and ‘cell differentiation’ (Figure 2). Together, these findings suggest that increased chromatin accessibility at promoter regions may contribute to altered expression of genes involved in, amongst others, cell migration-related processes, while decreased chromatin accessibility at promoters is associated with, amongst others, proliferation-related processes.

**Figure 2.**
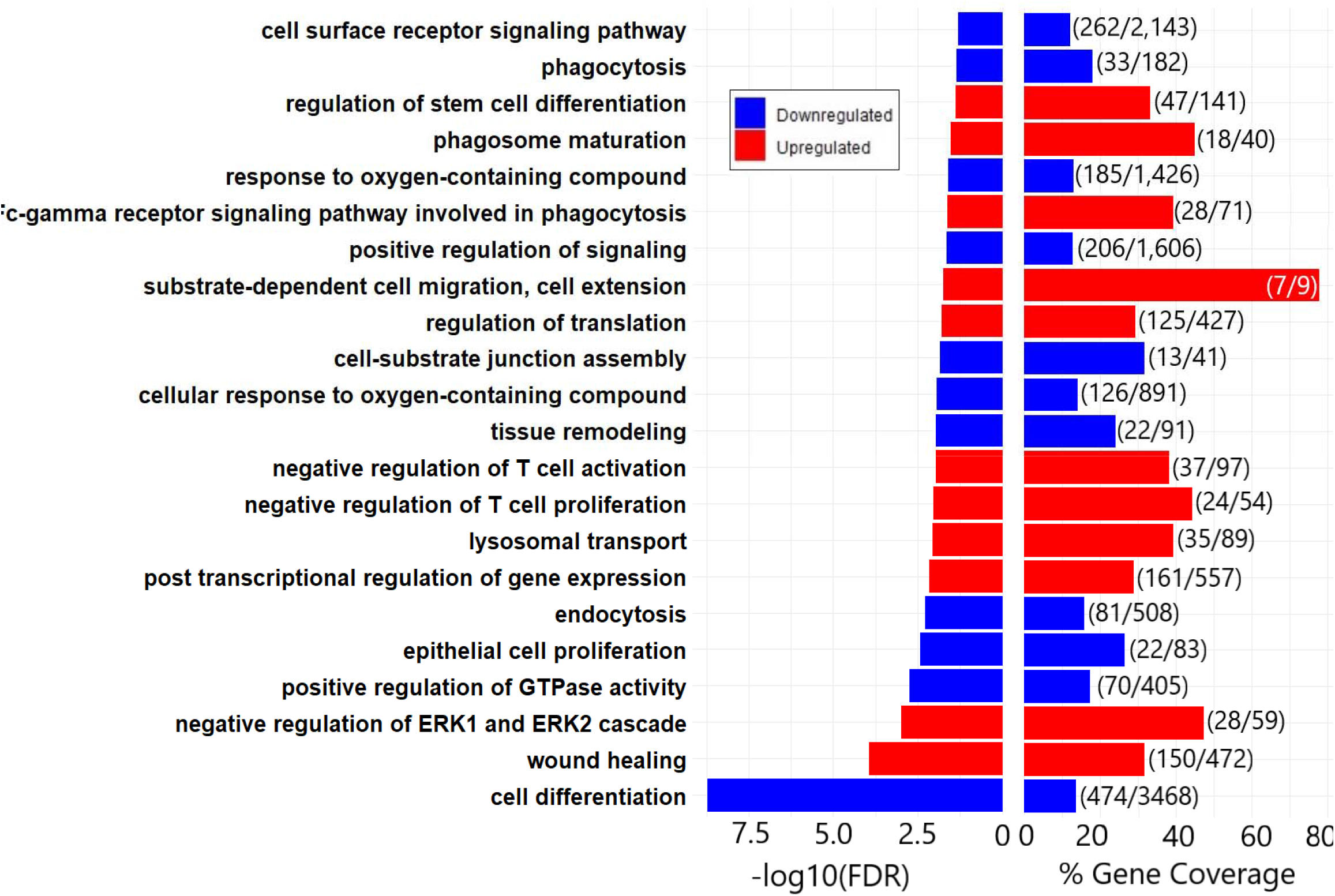
Functional analysis of genomic regions of significantly upregulated and downregulated DARs annotated to promot ers (0-3kb from annotated TSS), reveals increase in tumor progression processes. Bar plots depicting selected enriched up- and downregulated GO-BP terms among significantly upregulated and downregulated promoter (0-3 kb from annotated Transcription Start Site {TSS}). Promoter- Differentially Accessible Regions (DARs) were defined as statistically significant (FDR<0.05) ATAC-seq peaks of Nucleosome Free Regions (NFRs) within Open Chromatin Regions (OCRs), mapped to gene promoters. Upregulated (log_2_FC>0) DARs included 3,752 peaks, while downregulated (log_2_FC<0) DARs included 1,242 peaks. These DARs served as input for the rGREAT package in R programing. Downregulated (blue) processes correspond to those enriched among downregulated DARs (1,246 peaks), while upregulated (red) processes correspond to those enriched among upregulated DARs (3,570 peaks). FC, fold-change; FDR, false discovery rate; GO-BP, Gene Ontology biological Process; rGREAT, Genomic Regions Enrichment of Annotations Tool in R programing language.

### 3.3 Ectopic BORIS expression leads to increased promoter accessibility in conjunction with elevated gene expression at a subset of upregulated genes

To determine whether BORIS-mediated chromatin accessibility changes directly influence gene expression, we consequently aimed at identifying genes that displayed a concordat change in chromatin accessibility and gene expression in response to BORIS ectopic expression. The previously defined DEGs gene-list in the MM057 melanoma cells using RNA-seq, was achieved with the same comparison (BORpos) vs. controls (EVpos) (Janssen et al. 2020). The ATAC-Seq analysis was conducted concomitantly with RNA-Seq analysis, allowing for direct linkage of identified DEGs to DARs mapped to promoters of corresponding genes in the DEGs gene lists. The gene list of significantly (FDR<0.05) upregulated (log_2_FC>0) RNA-Seq-DEGs consisted of 1,307 genes, and the gene list based on ATAC-Seq-DARs mapped to promoters with increased accessibility (log_2_FC>0), consisted of 2,929 genes. The downregulated gene list (log_2_FC<0) of RNA-Seq-DEGs contained 935 genes and the ATAC-Seq-DARs mapped to promoters with decreased chromatin accessibility (log_2_FC<0), consisted of 881 genes. Taken together, these intersecting lists included 534 genes that showed significant differential accessibility of promoter region, coupled with significant mRNA differential expression in response to ectopic BORIS expression in melanoma cells (Figure 3). Concordance in directionality of expression and promoter accessibility changes was observed for all 534 genes. Most genes were upregulated (n=442; 83%, Figure 3A), while only 17% (n=92, Figure 3B) downregulated. Among the upregulated genes with overlapping DEGs and DARs directionality, were known cancer-related genes such as *PD-L1, GDF6, TNFA-IP2, IL1-β, CTLA4*, and *CDX2* (Figure 4).

**Figure 3.**
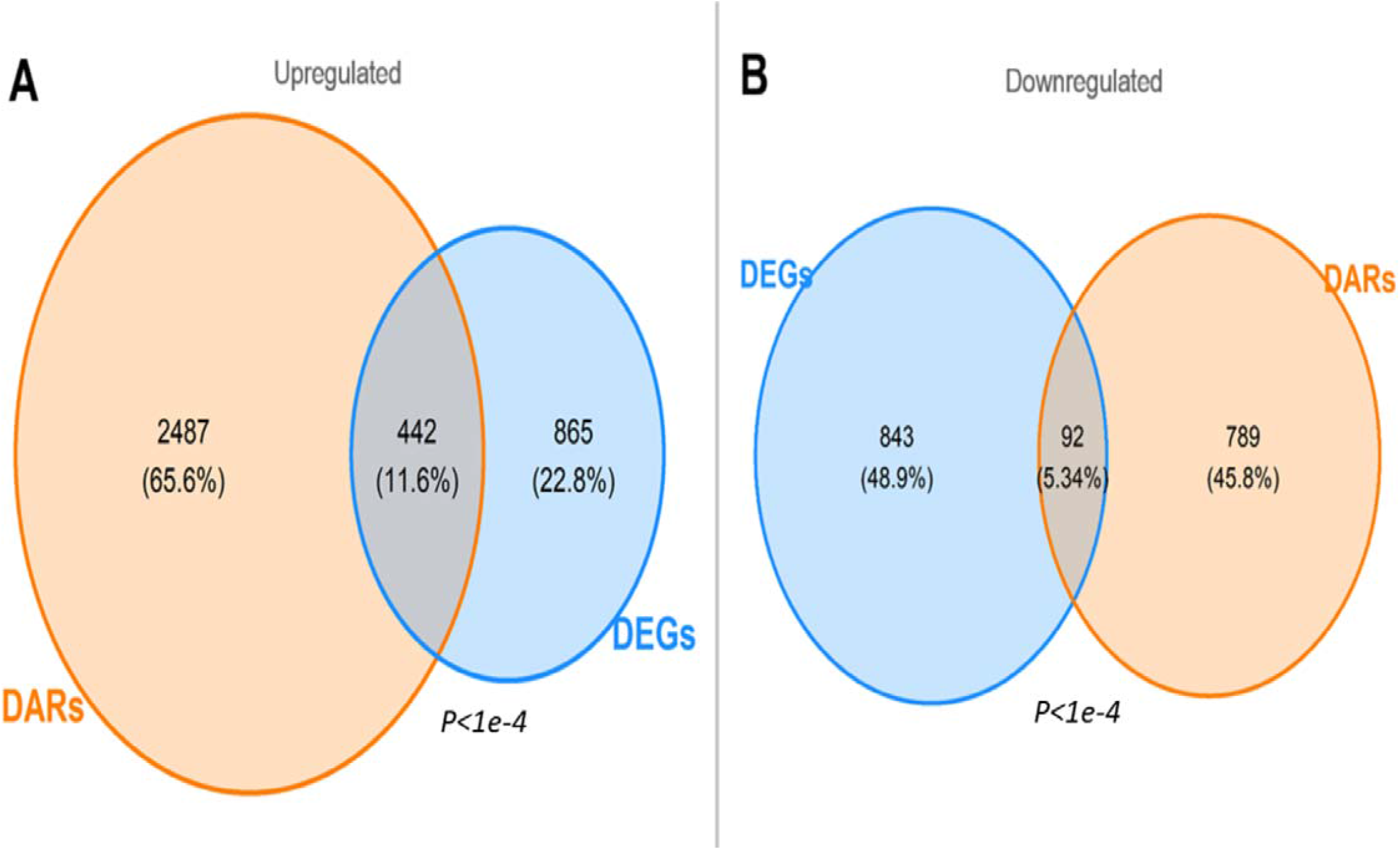
Intersects of ATAC-Seq DARS and RNA-seq DEGs yielded a DARs & DEGs gene-list. **(A, B)** Promoter-Differentially Accessible Regions (DARs) were defined as statistically significant (FDR<0.05 and |log_2_FC|>0) ATAC-seq peaks of Nucleosome Free Regions (NFRs) within Open Chromatin Regions (OCRs), that mapped to gene promoters (0-3kb from Transcription Start Site {TSS}), and further annotated to the corresponding gene. These DARs are a result of a differential analysis between EVpos versus BORpos samples. Thus, creating a promoters DARs based gene-list, that included 3,810 genes, 2,929 genes had significantly increased (log_2_FC>0) promoter accessibility, while 881 had significantly decreased (log_2_FC<0) promoter accessibility. The Differentially Expressed Genes (DEGs) gene-list was based on the same differential analysis of RNA-Seq expression data. The DEGs gene-list included 2,242 statistically significant (FDR<0.05) genes, out of them 1,307 were upregulated (log_2_FC>0) while only 935 were downregulated (log_2_FC<0). Intersection of the DARs gene-list and the DEGs gene-list resulted in 442 upregulated genes (log_2_FC>0) **(A)** and 92 downregulated genes (log_2_FC<0) **(B)**. The total number of genes defined in the DARs & DEGs gene-list was 534 genes. Both intersects were tested with the Chi-squared test and the results were statistically significant (p<1×10^−4^). ATAC-Seq, Assay for Transposase Accessible Chromatin followed by Sequencing; BORpos, BORIS-vector samples positive to doxycycline; Evpos, empty-vector samples positive to doxycycline; FC, fold-change; FDR, false discovery rate; RNA-Seq, RNA-Sequencing.

**Figure 4.**
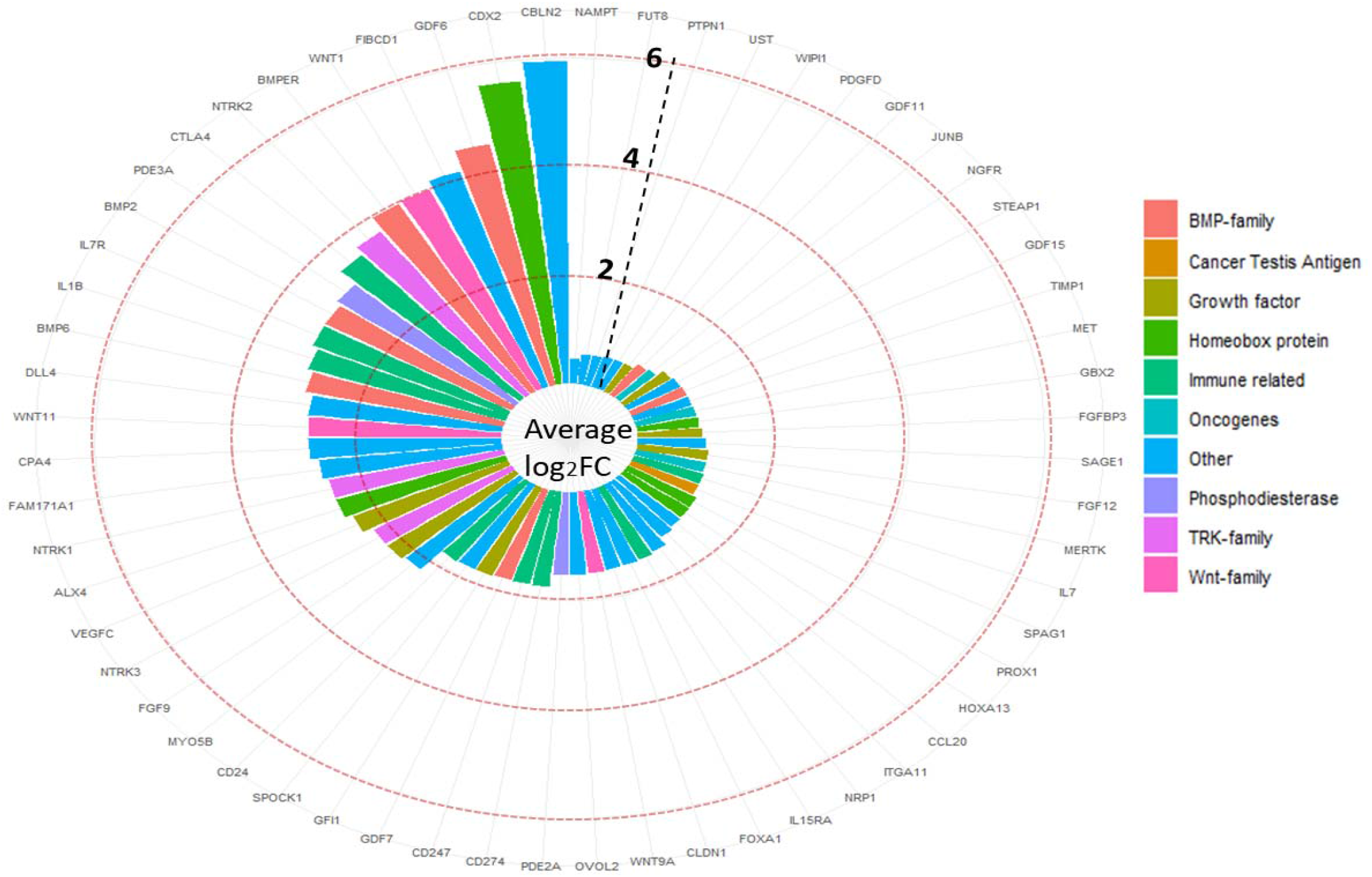
Circular bar plot of 60 tumor progressor genes, which were included in the upregulated intersect of DARs & DEGs gene-list. The genes were grouped by biological family or function, if available. Other group represents genes which did not belong to any particular pathway or could not be attributed to a protein family. The X-axis (outer circle characters) represents each gene’s name corresponding to its bar. Y-axis (inner circle bars) represents the average log_2_FC value from the two gene-lists: Differentially Accessible Regions (DARs) and Differentially Expressed Genes (DEGs). Each gene’s log_2_FC value from each list was summed and averaged, and this average is represented in the bar corresponding to the gene’s symbol. Light red dashed line at log_2_FC=2,4,6. FC, fold change. Light dashed black vertical line is for visual guidance.

**Figure 5.**
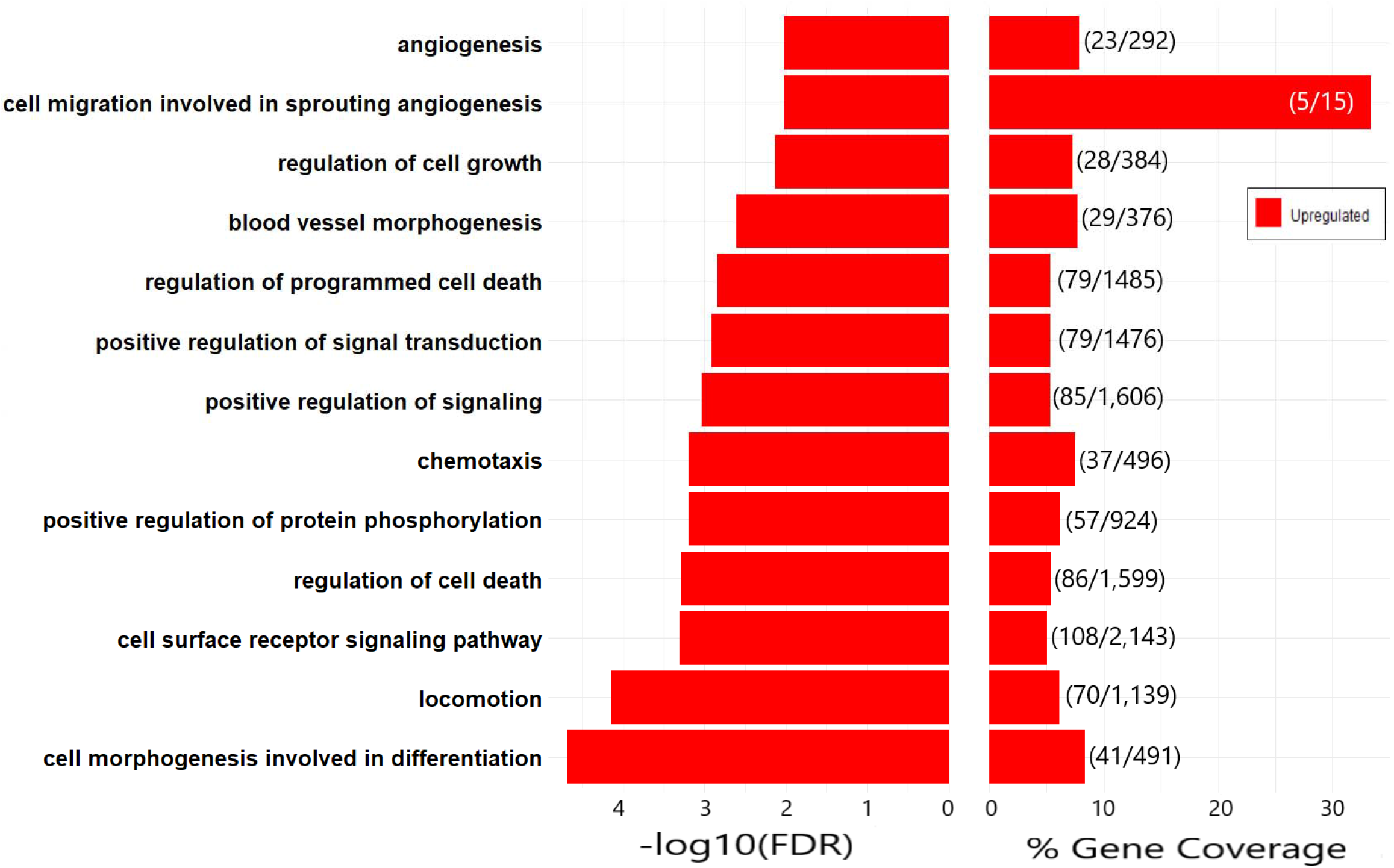
Functional analysis of promoter regions of genes defined as upregulated DARs & DEGs, identified processes in which ectopic BORIS expression is most likely to impact. Bar plots depict select enriched upregulated GO-BP terms among genomic regions mapped to promoters (0-3kb from Transcription Start Site {TSS}) of genes defined as upregulated Differentially Accessible Regions and Differentially Expressed Genes (DARs & DEGs). For the enrichment analysis, 554 promoters’ genomic regions were used as input for Genomic Regions Enrichment of Annotations Tool in R programing language (rGREAT). This subset of genes represents putative targets upregulated by BORIS, that met two criteria: their promoter region was identified with significant increased chromatin accessibility (i.e., DAR), and their mRNA transcript was identified as an upregulated DEG. The rGREAT package in R was used for enrichment analysis of the genomic regions. FDR, false discovery rate; GO-BP, Gene Ontology biological Process; mRNA, messenger RNA.

### 3.4 Enrichment analysis of genes with an ATAC-Seq peaks mapped to their promoter show high similarity when compared to the same RNA-Seq based analysis

To directly compare between RNA-seq-and ATAC-seq-identified differential genes and to characterize their associated enriched biological processes, an additional enrichment analysis was performed with the GSEA (Subramanian et al. 2005) tool. The analyses were preformed to test enrichment among the gene-sets representing GO-BP as well as melanoma invasive gene-set (Verfaillie et al. 2015). The analyses included all genes with an ATAC-seq peak mapped to its promoter (16,160 genes). The peaks were annotated to genes and pre-ranked by log_2_FC value of the results of a differential comparison between EVpos and BORpos samples, the same comparison that generated the DARs. The ATAC-Seq based gene list was created in a similar manner to the RNA-seq-based gene list, that served as input for the same GSEA analysis. Among all ATAC-Seq peaks mapped to promoters, 81.5% (13,170/16,160) showed increased chromatin accessibility (log_2_FC>0) and only 18.5% (2,990/16,160) showed decreased chromatin accessibility (log_2_FC<0). The ATAC-Seq-based GSEA analysis found significant (FDR<0.05) positive enrichment (NES>0) of 472 processes. In parallel, it revealed significant (FDR<0.05) negative enrichment (NES<0) for only 10 processes. A strong positive enrichment of EMT and cell migration processes was observed, as well as the melanoma-specific invasive gene signature (Figure 6). These data agree with the RNA-seq analysis and further support the suggestion that ectopic BORIS expression promotes an invasive gene signature and phenotype in melanoma cells by altering chromatin accessibility.

**Figure 6.**
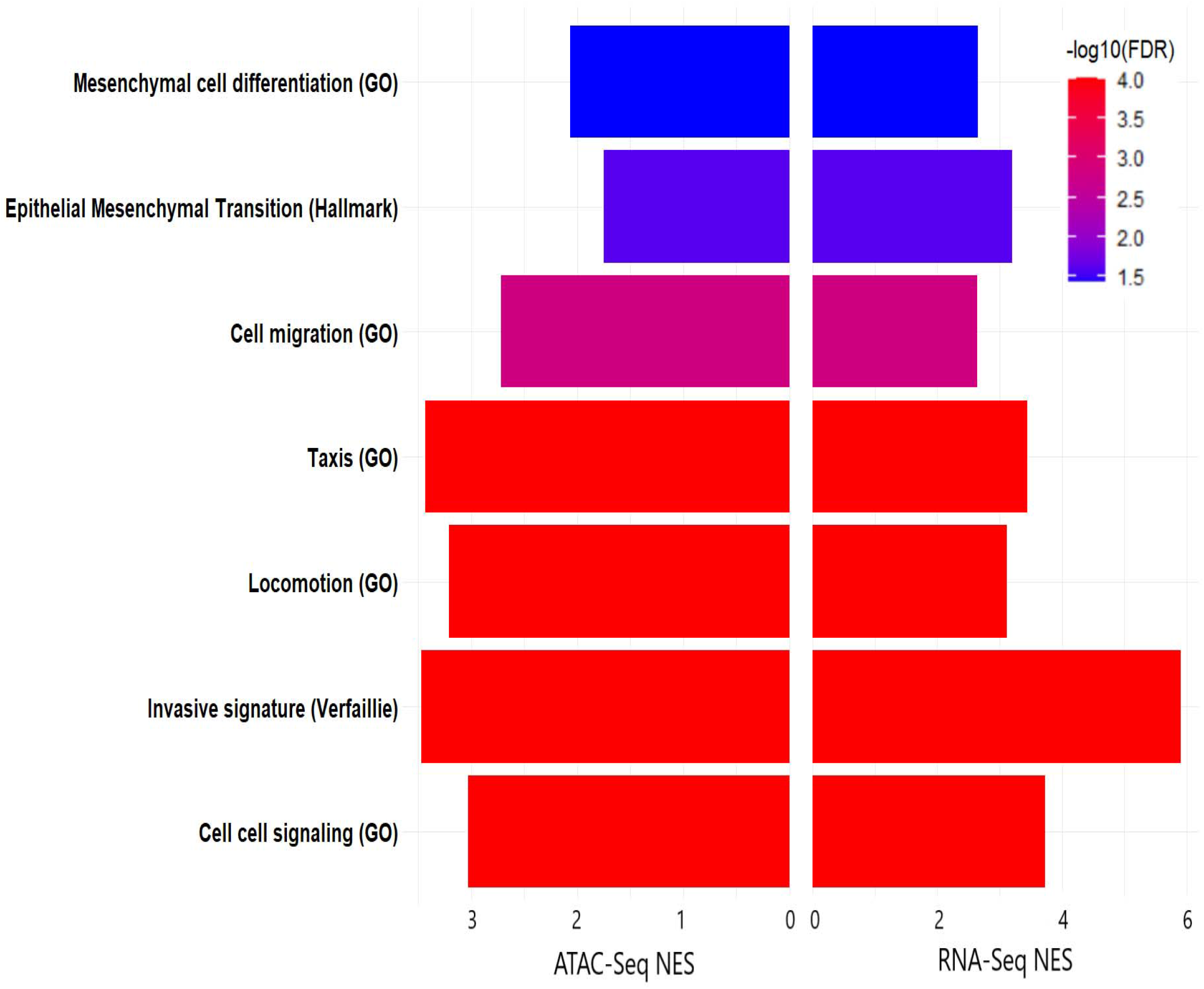
GSEA of ATAC-Seq as compared to GSEA of RNA-Seq, revealed similar enriched processes. In both experiments (ATAC-Seq and RNA-Seq), gene-lists which served as input for Gene Set Enrichment Analysis (GSEA), were ranked according to a differential analysis comparing MM057 melanoma cells expressing a Dox-inducible vector, that either contained the gene BORIS (BORpos) or an Empty-Vector (EVpos). Normalized Enrichment Score (NES) results of GSEA are presented in bar plots representing processes in which similar enrichment was found in both analyses. The selected processes of GO-BP, hallmark and EMT gene sets from the MSigDB and the invasive gene signature of Verfaillie et al.^7^ are presented. On the left side of the plot, GSEA NES results based on ATAC-Seq peaks mapped to promoters (0-3kb) and further annotated to their nearest gene, are presented. This gene list summed to a total of 16,160 genes. The gene list was ranked according to the value of log_2_FC in promoter accessibility of the gene. On the left side of the plot, GSEA NES results of RNA-seq data gene-list which is described in Fig. 3 (A-C), are presented. The GSEA-pre-ranked algorithm (version 4.1.0) was used. NES>2 is considered significant enrichment of the tested processes. The bar color represents the processes’ –log10(FDR) value, from less significant (blue) to more significant (red) processes. The processes are ordered by statistical significance, with the most significant are at the bottom of the plot. ATAC-Seq peaks mapped to promoters (0-3 kb) were annotated to their nearest gene with ChIPseeker in R programing. ATAC-Seq, Assay for Transposase Accessible Chromatin followed by Sequencing; BORpos, BORIS-vector samples positive to doxycycline; Dox, doxycycline; EMT, Epithelial-Mesenchymal Transition; Evpos, empty-vector samples positive to doxycycline; FC, fold-change; FDR, false discovery rate; GO-BP, Gene Ontology biological Process; MSigDB, Molecular Signatures Database.

### 3.5 Integrated ATAC-seq and RNA-seq analysis identifies differential transcription factor activity, including of cancer-related TFs, upon ectopic BORIS expression

To gain a deeper insight into BORIS-mediated transcriptional regulation in melanoma cells, an assessment of differential TF activity was conducted in EVpos compared to BORpos cells using diffTF (Berest et al. 2019; Rasmussen et al. 2019). Briefly, diffTF is a novel multi-platform software framework that estimates differential TF activity based on differences in chromatin exposure around TF DNA-binding sites, as captured by ATAC-Seq. Integration of the differential chromatin accessibility with RNA-seq-determined gene expression, yields a profile of TFs that are predicted to activate or repress transcription. Each TF classified as an activator/repressor displays increased/reduced accessibility around its DNA-binding motif as captured by ATAC-Seq, and upregulation/downregulation of its target genes as captured by RNA-Seq. This integrated analysis revealed ectopic BORIS expression differentially impacted 32 activators and 38 repressors TFs, in a statistically significant manner (FDR<0.05). To further narrow the list of TFs, those that were suspected to be differentially activated or repressed by dox, as determined by a comparison of EVneg vs EVpos, were rejected. All TFs classified as activators or repressors, with a relaxed statistical cutoff (FDR<0.1), in the EVneg vs EVpos comparison, were compared to the significant TFs identified in the comparison of EVpos vs BORpos. The overlap between the two comparisons included 11 activators and 13 repressors TFs, which were rejected. Closer examination of the results found several TFs that bind more than one motif. TFs with a double motif that were classified as activators were: *MAZ* and *SP1*, while TFs classified as repressors included *NF1A, TP73*, and *SOX10*. Thus, the final list of TFs differentially regulated by ectopic BORIS expression was further reduced to 19 activators and 22 repressors (Figure 7). Activators included various TFs associated with tumor progression, such as *SP1, STAT6, HOXA1, PBX3, NFYB, NFYC*, and *TEAD4*. Notably, *TEAD4*, the top activator TF reported by diffTF (Figure 7a), was previously shown to act as an invasiveness-linked TF in melanoma (Zhang et al. 2018; Verfaillie et al. 2015). Conversely, TFs known to act as tumor suppressors in melanoma, including *TP53* (Bardeesy et al. 2001) and *TFAP2A* (Seberg et al. 2017a), are preferentially repressed by BORIS (Figure 7b). In addition, SOX10 (Shakhova et al. 2012), a proliferation-linked TFs in melanoma, was identified as a repressor TFs (Figure 7B), supporting the previously observed BORIS-mediated reduction in proliferation. Together, diffTF identified TFs that are likely differentially regulated by BORIS and suggest that BORIS-induced changes in gene expression could be mediated, at least in part, by altered activity of cancer-related TFs.

**Figure 7.**
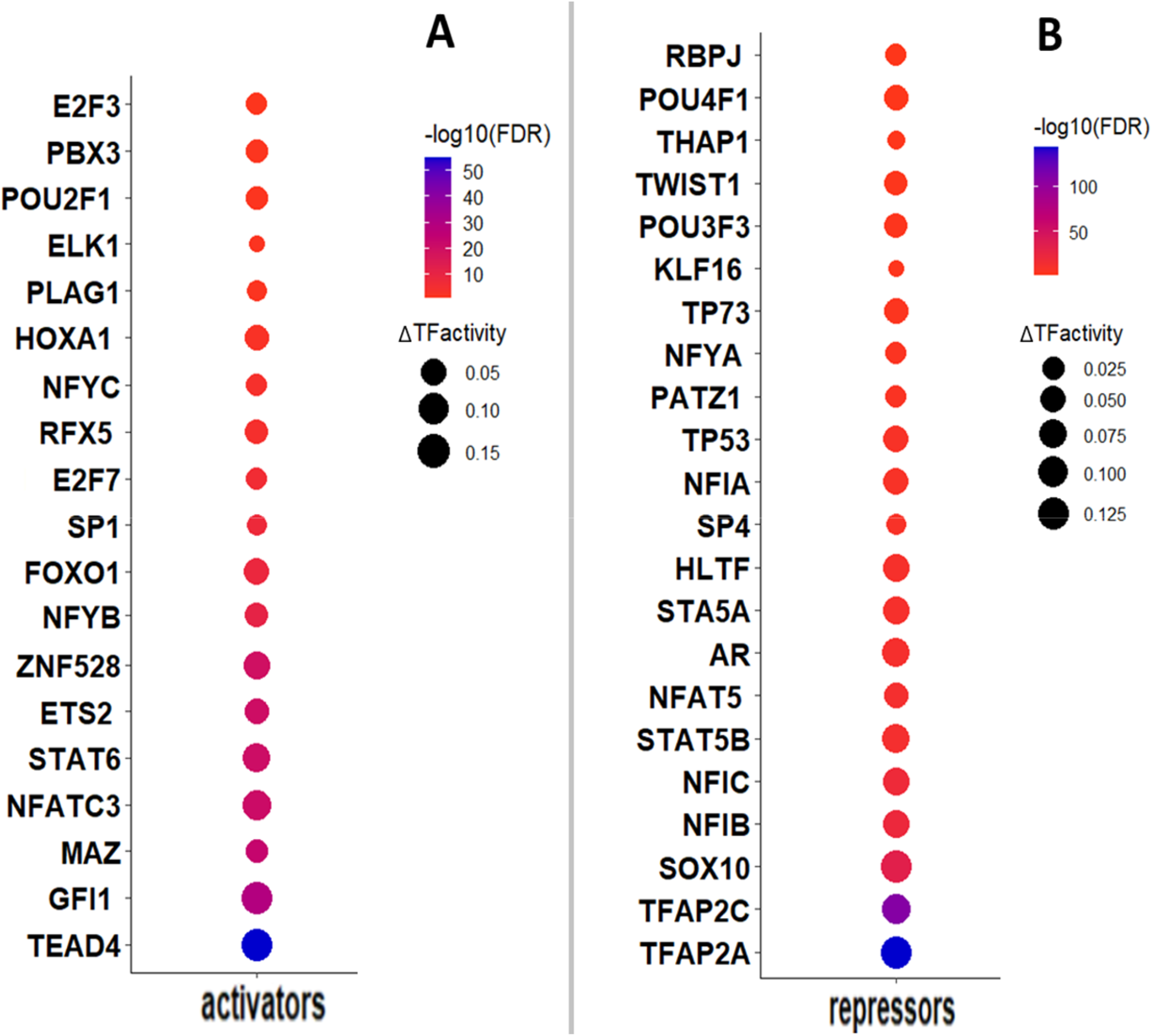
Differential activity analysis of transcription factors (TFs) reveals tumorigenesis TFs as activators and tumor suppressors as repressors, upon ectopic BORIS expression in melanoma cells. Results of the diffTF software framework in Transcription Factor (TF) classification mode. Displayed are TFs whose DNA-binding motifs showed statistically significant differential activity as activators (A) or repressors (B) in MM057 melanoma cell-line expressing an Empty Vector (Evpos) versus those expressing a BORIS vector (BORpos). TFs were classified as activators or repressors based on increased or decreased ATAC-Seq chromatin accessibility of 100 bp around the DNA-binding motif and the upregulation or downregulation of their target genes as captured by RNA-Seq. The dot color represents the TF’s –log_10_(FDR) value, with the TFs ordered by statistical significance, from less significant (red) to more significant (blue) TFs. The dot size represents the magnitude of the change in activity (delta, ΔTF), with bigger size dot signifying greater change in differential activity. TF DNA-binding motifs were defined by PWM database HOCOMOCO. ATAC-Seq, Assay for Transposase Accessible Chromatin followed by Sequencing; BORpos, BORIS-vector samples positive to doxycycline; bp, base pairs; Evpos, empty-vector samples positive to doxycycline; FDR, false discovery rate; PWM, Position Weight Matrix; TFs, transcription factors

## 4 DISCUSSION

BORIS is a cancer testis antigen involved in various types of cancer, including melanoma, through still largely unknown mechanisms (Loukinov et al. 2002; Kholmanskikh et al. 2008; Soltanian and Dehghani 2018). In a previous study, we showed that BORIS contributes to phenotype switching from a proliferative to an invasive state by altering the transcriptome of melanoma cells (Janssen et al. 2020). Here, we set out to gain further insight into the mechanisms through which BORIS expression induces transcriptional changes, focusing on chromatin accessibility. Using the MM057 melanoma cell line, which harbors a proliferative transcriptome (Verfaillie et al. 2015; Wouters et al. 2020; Janssen et al. 2020), we show that dox-inducible BORIS expression leads to altered chromatin accessibility. Most differential changes in chromatin accessibility (i.e., DARs) involve gains in accessibility.

We investigate differential changes in promoter accessibility (defined as 0-3 kb from annotated TSS). Integration of the chromatin accessibility data with gene expression data reveals that for a subset of putative BORIS target genes, gain/loss in promoter accessibility is accompanied by corresponding up/down regulation of gene expression (i.e., the DARs & DEGs intersect). Among all identified genes with an ATAC-Seq peak mapped to their promoter we find enrichment for the melanoma-specific invasive gene signature and invasion-related processes, like EMT. Notably, similar enriched processes and gene signatures are found when comparing the ATAC-Seq gene enrichment results to previously obtained RNA-Seq based enrichment analysis from the same cell culture model system. This indicates that the observed chromatin accessibility changes at gene promoters are concordant with transcript expression changes, and further support BORIS’ contribution to transcriptional phenotype switching.

It has been suggested that BORIS functions as a transcriptional regulator (Hong et al. 2005; Liu et al. 2017; Bergmaier et al. 2018). In agreement, here we show that BORIS-induced changes in chromatin accessibility likely contribute to the altered transcriptional landscape of the MM057 melanoma cell line. We also find that ectopic BORIS expression in this cell line with a proliferative transcriptional state induces aberrant chromatin accessibility and upregulated expression of some cancer-related genes, that are involved in invasive cell behavior and malignant progression.

PD-L1 (CD274) and CTLA4 genes are both differentially upregulated by BORIS and have increased chromatin accessibility at their promoter regions, suggesting that chromatin remodeling related to BORIS is a possible modulator of the regulatory T cells microenvironment.

Interestingly, the key melanoma metastasis driver gene *Growth Differentiation Factor 6* (*GDF6)* is ranked at the top of the DARs & DEGs gene list. GDF6 is a ligand of the Bone morphogenetic proteins (BMPs) signaling pathway, required for tumor maintenance, and was pointed as a key factor in melanoma tumor progression and the formation of metastasis (Venkatesan et al. 2018). *GDF6* has been reported as amplified and transcriptionally upregulated in melanoma, where it helps to maintain a trunk neural crest gene signature. Additionally, GDF6 represses MITF and the pro-apoptotic factor SOX9, therefore preventing differentiation, inhibiting cell death, and supporting tumor growth (Venkatesan et al. 2018).

Additional members of the GDF family were identified as differentially regulated by BORIS, including GDF7, GDF11 and GDF15. GDF15 is a serum marker for metastases in uveal melanoma patients (Suesskind et al. 2012) and is linked to PD-L1 expression regulation in glioblastoma (Peng et al. 2019). GDF11 upregulation is linked to shorter overall-survival for uveal melanoma (Liu et al. 2019). Of note, BMPs are ligands of the transforming growth factor beta (TGF-β) superfamily, and BORIS expression activates TGF-β in both neuroblastoma (Makani et al. 2021) and melanoma as we revealed previously in the same MM057 melanoma cell line model (Janssen et al. 2020). In our current study we have linked TGF-β targets such as NGFR and JUNB, which are regulators of melanoma phenotype switching (Verfaillie et al. 2015; Restivo et al. 2017), to ectopic BORIS expression in the MM057 cell line, as those genes were identified among the DARs & DEGs intersected gene list.

Examining the small downregulated DARs & DEGs intersect gene list (Figure 3b), revealed a noticeable tumor suppressors among them, the gene *missing in metastasis protein (MTSS1)*, an invasion preventor gene (Vadakekolathu et al. 2018). Finding *MTSS1* activity as repressed by BORIS may explain how it may support metastasis formation.

To explore the impact of ectopic BORIS expression on transcriptional regulation in melanoma cells in more depth, TFs activity was assessed using diffTF (Berest et al. 2019). DiffTF is a multi-platform tool that combines ATAC-Seq and RNA-Seq data to identify TFs that are differentially activated or repressed between two conditions. Here, diffTF was used for the comparison of control melanoma cells (EVpos) to melanoma cells expressing ectopic BORIS (BORpos), to reveal aberrantly activated or repressed TFs upon ectopic BORIS expression. TEAD4 is the most significantly differentially activated TF. In line with this finding, a previous study identified the AP-1/TEAD families of TFs as master regulators of the melanoma-specific invasive gene network and established a causal link between TEADs and melanoma cell invasion (Verfaillie et al. 2015), highlighting TEAD4 as a potential candidate TF through which BORIS may promote melanoma phenotype switching. The second most significant TF is GFI1, a zinc finger TF which was identified here as aberrantly activated. Even though its role in melanoma remains unclear, GFI1 can act as a tumor promoter or suppressor depending on tumor type and has been shown to bear a wide range of cellular activities including inhibition of cell proliferation (Ashour et al. 2020). Future studies on GFI1 in melanoma will be important to obtain a better understanding of a potential role for GFI1 in melanoma phenotype switching. Additionally, oncogenic Tyrosine Kinase ELK1 was found as activated by BORIS. ELK1 has been linked to the AP-1 TFs network, as they coordinately increase expression in late stage 4 melanoma, implying these TFs may contribute to melanoma progression (Singh et al. 2020).

When turning to analyze the differentially repressed TFs, we first examined the melanocytic lineage specific TF MITF and no significant differential change in its activity was observed.

However, MITF activity has been suggested to be dependent on the TFs acting alongside it, as different transcriptional combinations result in the activation of proliferation or invasion targets (Seberg et al. 2017a). Here, two of the MITF’s main pro-proliferative collaborators, TFAP2A and SOX10 were classified as the most and the third-most significantly differentially repressed TFs (Figure 7B). SOX10 is a well-known regulator of the melanoma-specific proliferative gene network and its protein is widely used as a diagnostic biomarker of the melanocyte lineage (Verfaillie et al. 2015) and TFAP2A exhibits decreased expression in the advanced stages of melanoma (Seberg et al. 2017b). These findings suggest BORIS might shifts MITF activity by repressing its pro-proliferation collaborators, while activating pro-invasive collaborators.

Other proliferative related TFs in melanoma, such as STAT5A, STAT5B (Hassel et al. 2008; Morcinek et al. 2002), AR (Ma et al. 2021), and POU4F1 (Hohenauer et al. 2013) are classified with repressed activity. In addition, almost all members of the Nuclear Factor one (NFI) family; NFIA, NFIB and NFIC, were classified as repressed TFs. This family of TFs interact with chromatin, which results in increased chromatin accessibility and gene expression (Pjanic et al. 2013). Their role is context-dependent across different cancer types, with some being oncogenes and others tumor suppressors (Fane et al. 2017a; Becker-Santos et al. 2017). We propose two options regarding NFI-family activity in the context of ectopic BORIS expression in melanoma: either BORIS overrides their oncogenic activity, especially NFIB, which has been shown capabilities in driving invasive phenotypic in melanoma (Fane et al. 2017b), by repressing them, in favor of a BORIS-invasiveness genetic signature, or they function here as tumor suppressors, hence their repressed activity.

These observations suggest that ectopic BORIS expression facilitates de-repression of invasion-associated genes via loss of chromatin accessibility at the promoters, and in turn enhances transcriptional repression of NFIA, NFIB and NFIC. If re-expression of NFI family members or loss of TEAD4 in melanoma cells with ectopic BORIS expression can reverse or prevent the melanoma phenotype switching observed upon ectopic BORIS expression warrants further investigation.

A plausible mechanism by which BORIS may alter promoters accessibility is by activating the two non-specific DNA interacting subunits NFYB/NFYC of the Nuclear Transcription Factor Y (NF-Y) complex (Oldfield et al. 2019), whilst repressing its DNA sequence specific subunit NFYA (Nardini et al. 2013; Huber et al. 2012), thus maintaining open chromatin promoters accessibility at non-specific locations across the genome, including in cancer-related genes identified here.

Altogether, our integration of ATAC-Seq and RNA-Seq data demonstrates that BORIS mediates transcriptional reprogramming in melanoma cells by altering chromatin accessibility. These differential changes, mainly gain of accessibility at promoters, are associated with transcriptional upregulation of genes involved in tumor progression processes, and the aberrant activation of invasive signature oncogenic transcription factors, while reduced promoter accessibility and downregulated genes, are associated with repressed activity of proliferative signature factors and tumor suppressors. Our findings demonstrate BORIS chromatin remodeler capabilities of shifting the cellular transcription landscape of proliferative melanoma cells towards a pro-invasive genetic signature.

## ACKNOWLEDGEMENTS

This work was partially funded by the Israel Cancer Research Fund. S.M.J. was supported by the Banque National Fellowship, the Dr. Victor K.S. Lui Fellowship and the CIHR/FRSQ training grant in cancer research (FRN53888) of the McGill Integrated Cancer Research Training Program. A.I.P. was supported by the Banque National Fellowship.

We thank Dr Ghanem Ghanem for generously providing the MM57 cell line.

---

**Figure S1.**
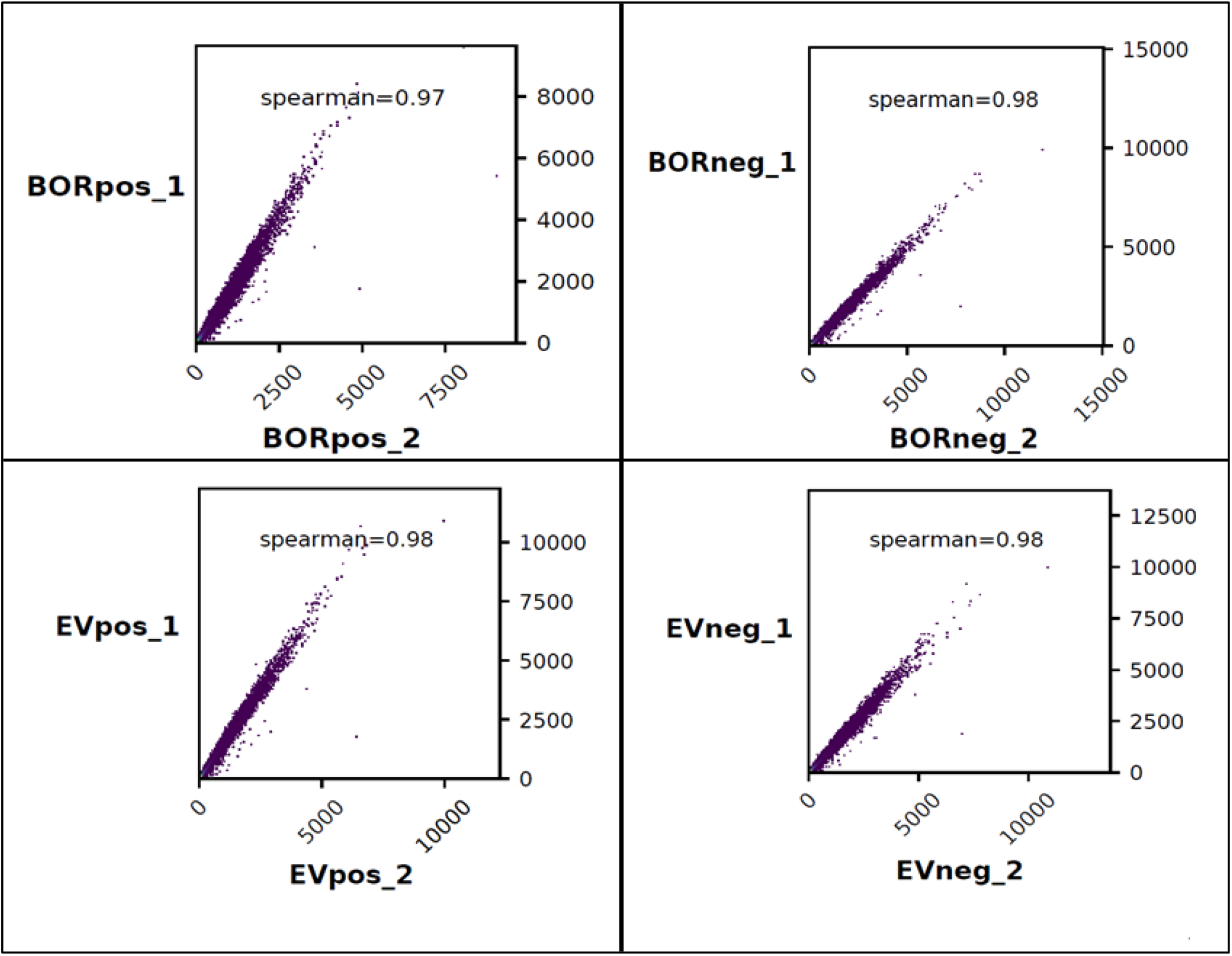
Spearman correlation between replicates based on ATAC-Seq BAM files. The X and Y axis represent the number of fragments in each repeat. Plots were generated with deepTools2 plotCorrelation. EVneg: empty-vector samples negative to doxycycline, EVpos, empty-vector samples positive to doxycycline; BORneg, BORIS-vector samples negative to doxycycline; BORpo, BORIS-vector samples positive to doxycycline; 1=biological repeat number 1, 2= biological repeat number 2.

**Supplement 2.**
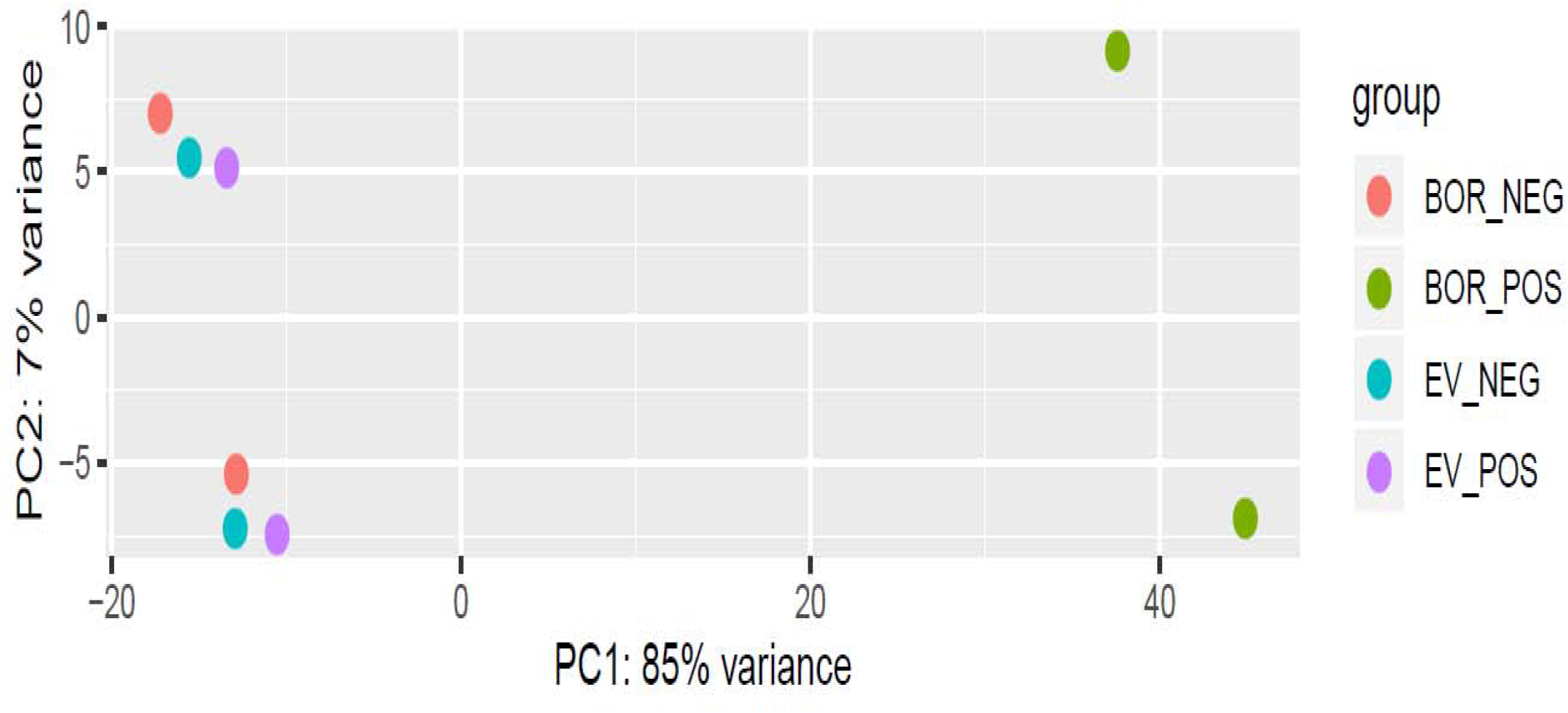
Principal Component Analysis (PCA) of the four conditions of the ATAC-seq samples. The PCA was plotted with DESeq2^194^ in R software. EV_NEG, empty-vector samples negative to doxycycline; EV_POS, empty-vector samples positive to doxycycline; BOR_NEG, BORIS-vector samples negative to doxycycline; BOR_POS, BORIS-vector samples positive to doxycycline; PC1, principal component 1; PC2, principal component 2

**Supplement 3.**
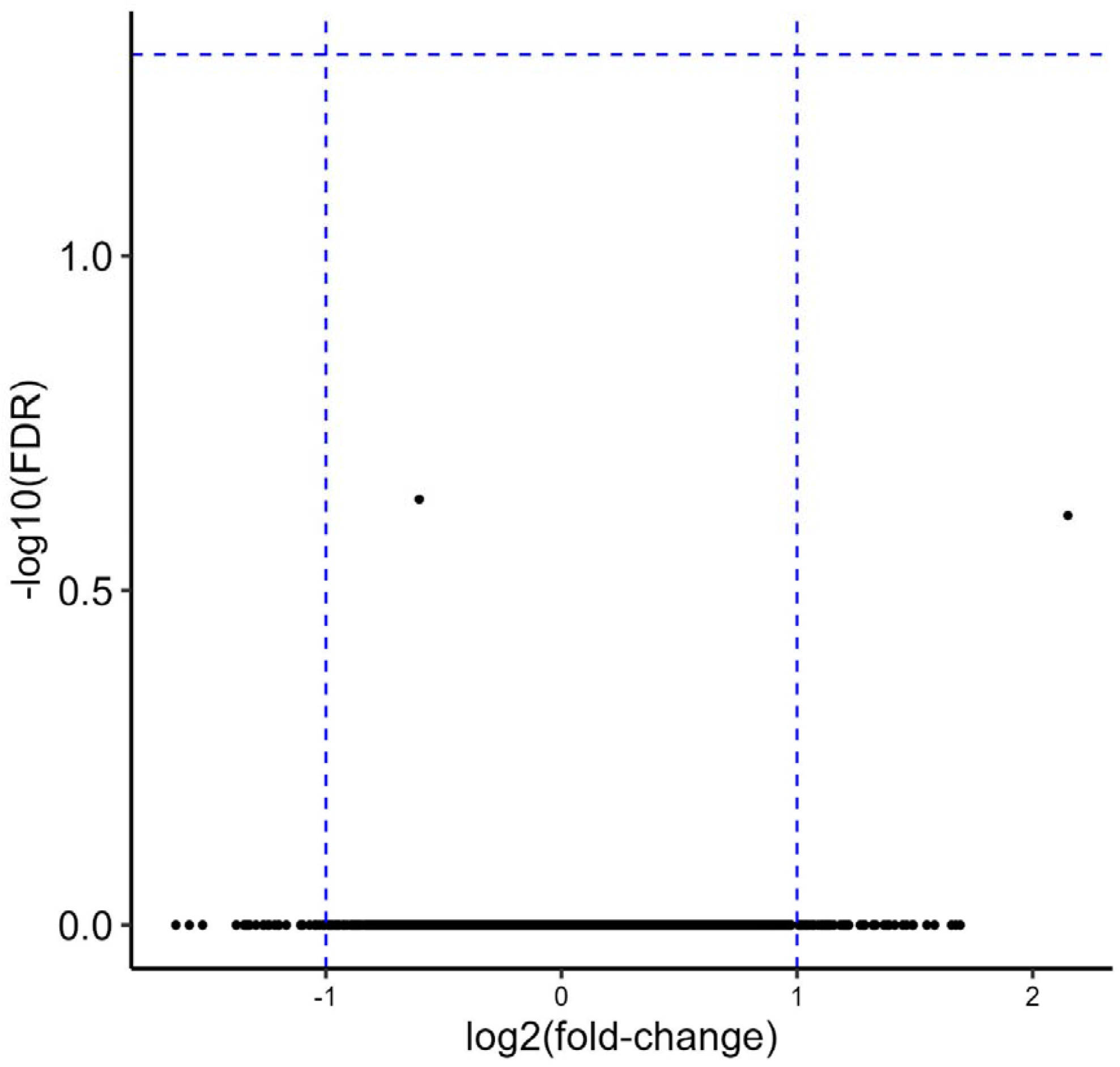
Empty-Vector (EV) expression in melanoma cells leads to no significant differentially accessible open chromatin. Volcano plot of ATAC-seq differential peaks comparison between EVneg and EVRpos samples. Differentially accessible regions (DARs) corresponded to 18,897 genes presented in the plot. None were statistically significantly (FDR<0.05) up-or downregulated in the EVpos samples compared to the EVneg samples. log_2_fold-change represents result of a differential analysis between Evneg to EVpos samples. Blue dashed lines: X-axis at X=1, X=-1. Y-axis, Y=1.30102999566 (FDR=0.05). For the differential analysis, Bioconductor DESeq2 was used. ATAC-Seq, Assay for Transposase Accessible Chromatin followed by Sequencing; Evpos, empty-vector samples positive to doxycycline; Evneg, empty-vector samples negative to doxycycline. FDR, false discovery rate

**Supplement 4.**
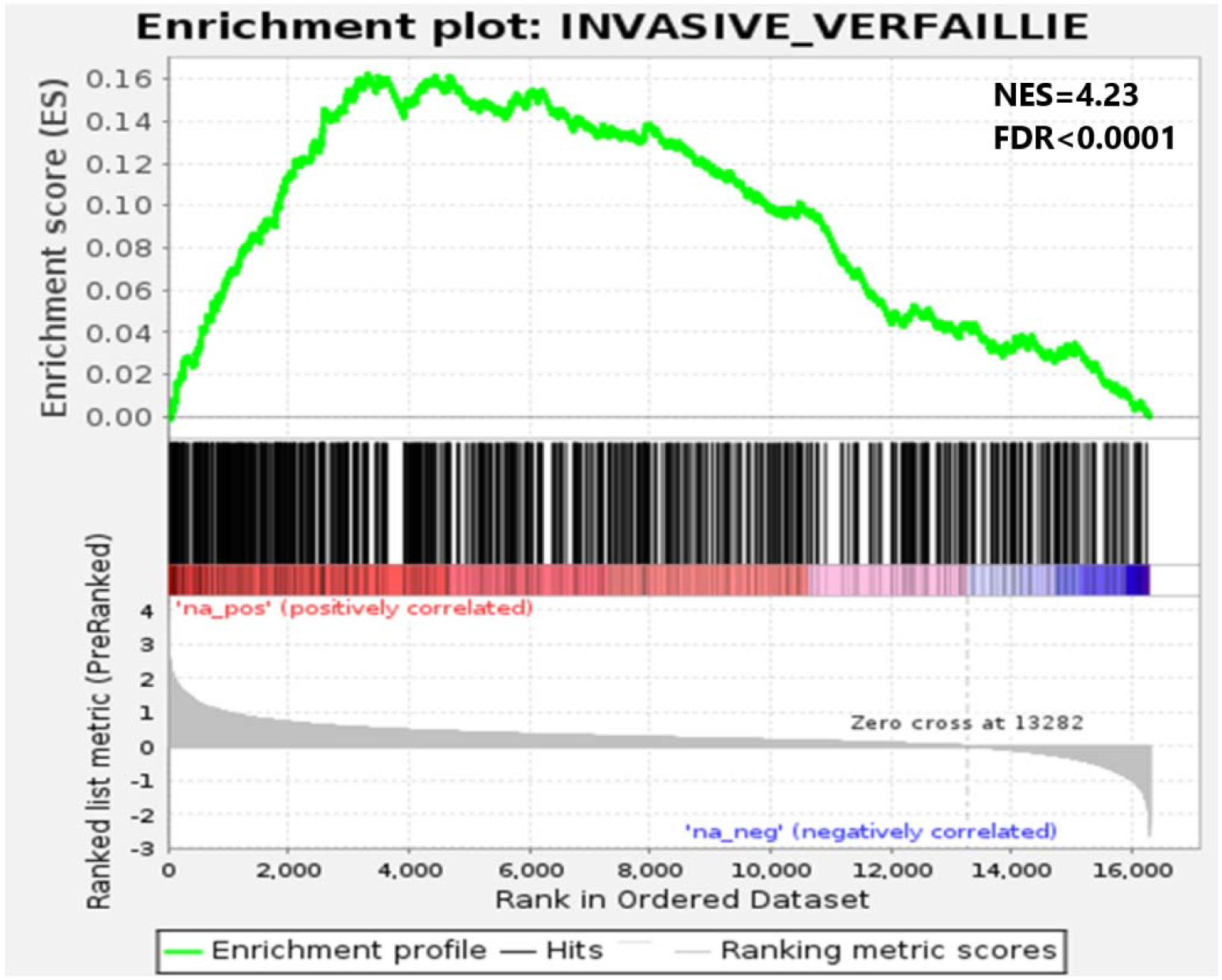
Verfaille invasive gene set signature barcode plots of all ATAC-Seq peaks identified. All 16,161 genes, which their promoter (0-3kb) region had ATAC-Seq peaks who mapped to it, served as input for the Gene Set Enrichment Analysis (GSEA). Genes were ranked according to a differential analysis comparing MM057 melanoma cell line expressing a DOX inducible vector, that either contained the gene BORIS (BORpos) or an empty-vector (EVpos). The genes were ranked according to their log_2_FC and served as input for GSEA-pre-ranked algorithm (version 4.1.0). NES, Normalized Enrichment Score; FDR, False Discovery Rate.

**Supplement 6.**
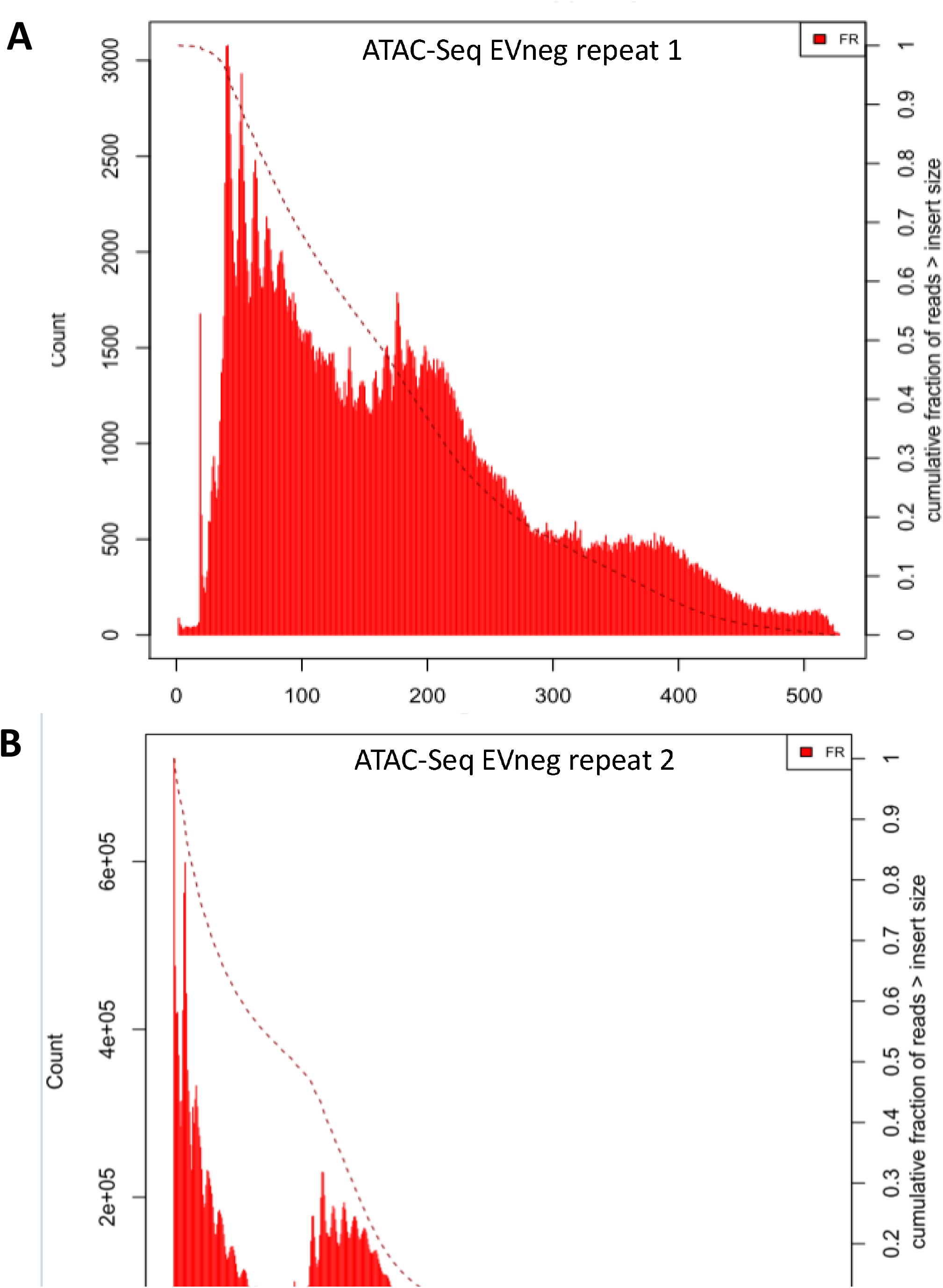
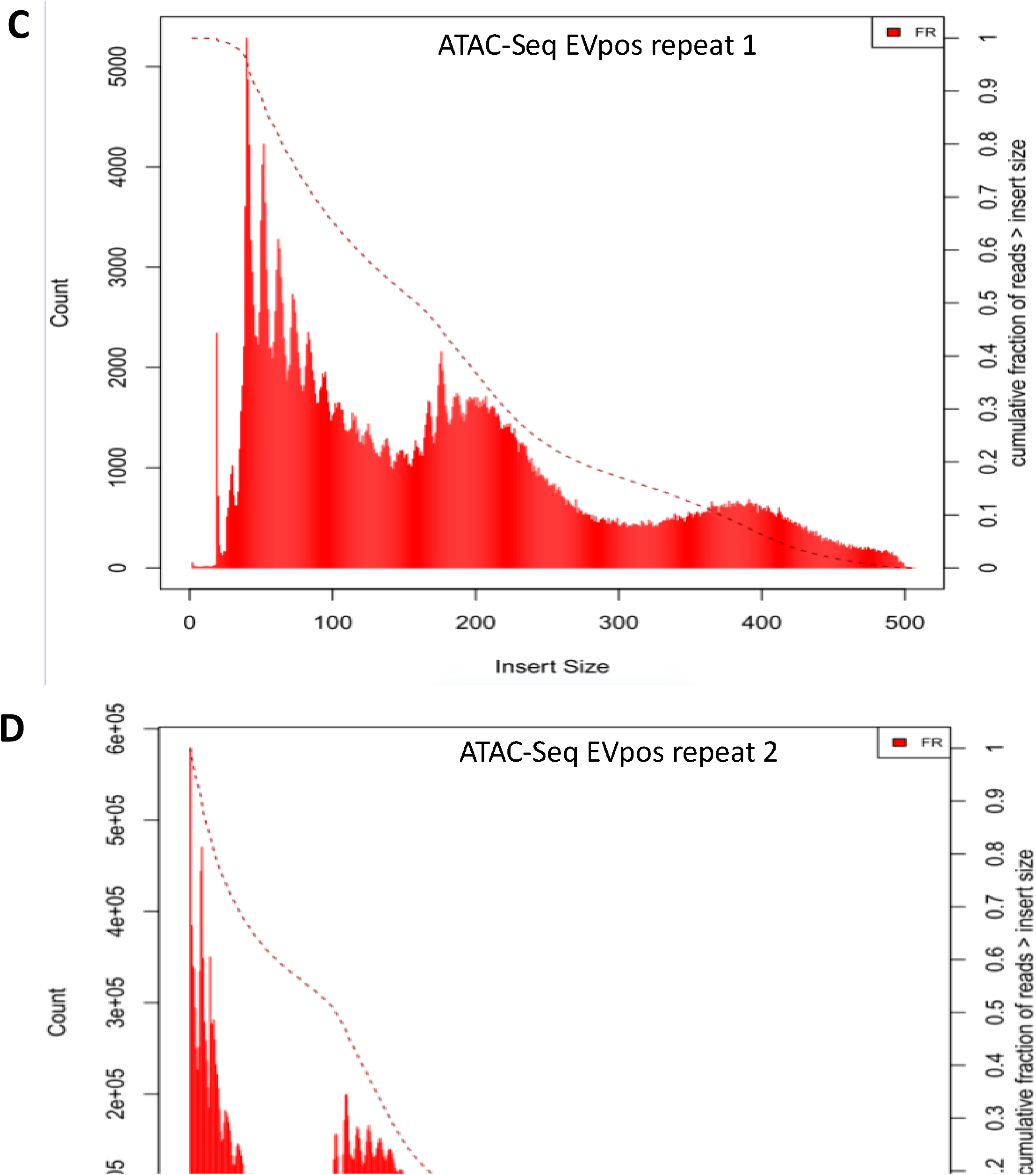
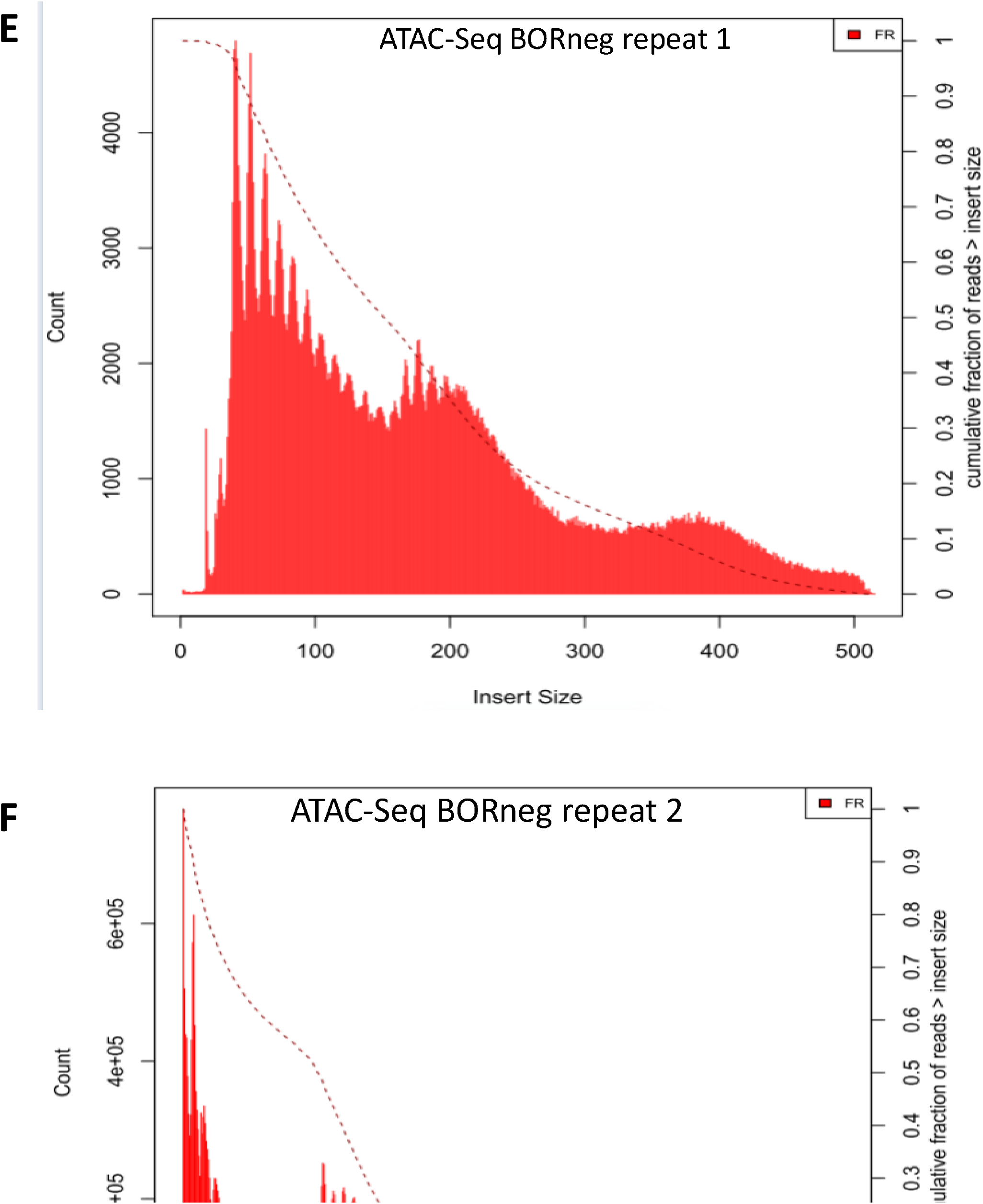
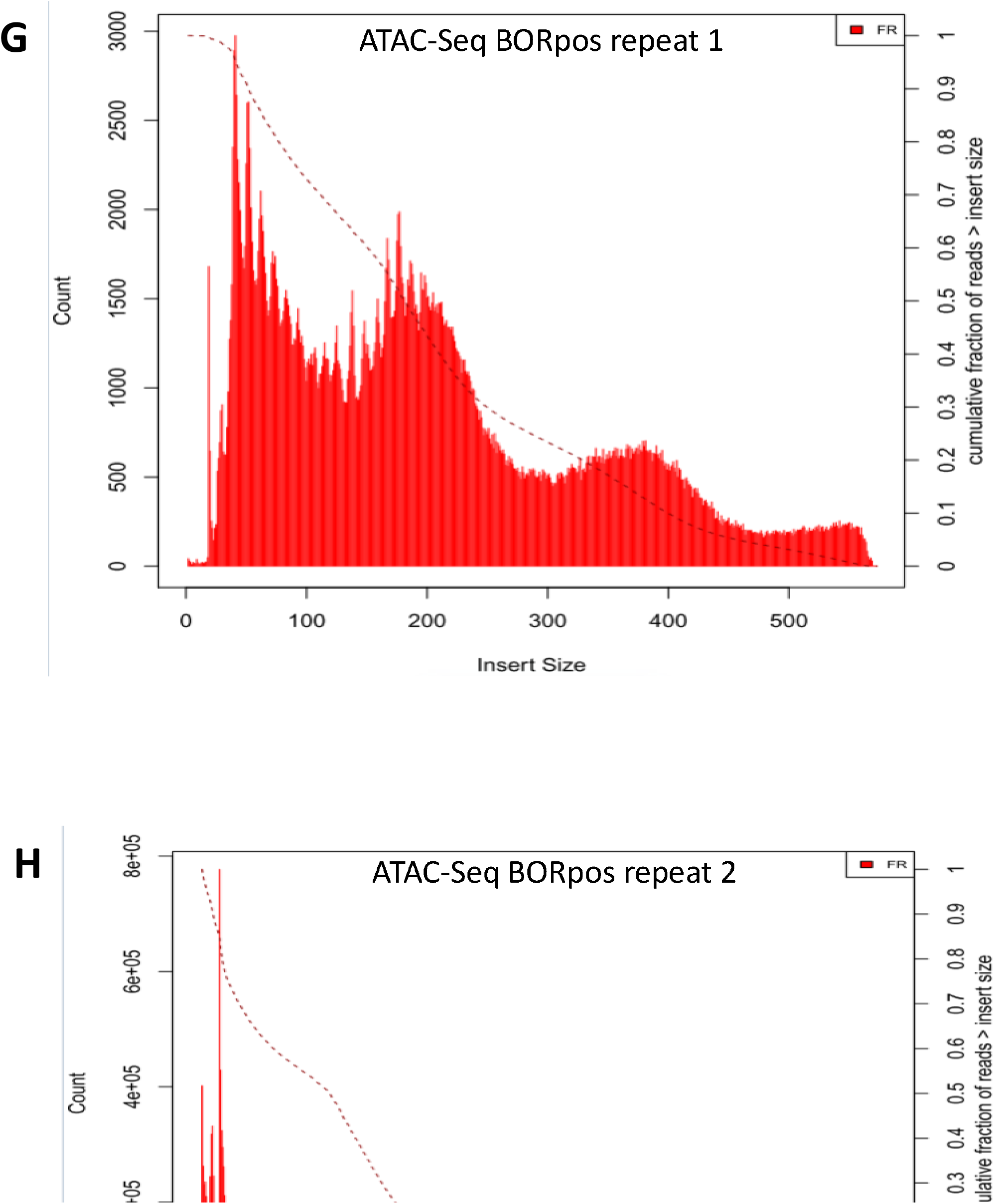
Distribution of ATAC-Seq fragment sizes, over the four experimental conditions EVneg,EVpos,BORneg,BORpos. Results as produced by PicardTools: CollectInsertSizeMetrics. The X-axis represents the number of fragments. The Y-axis represent the fragment size in base-pairs (bp). Evpos, empty-vector samples positive to doxycycline; BORpos, BORIS-vector samples positive to doxycycline; Evneg, empty-vector samples negative to doxycycline; BORneg, BORIS-vector samples negative to doxycycline

**Supplement 7.**
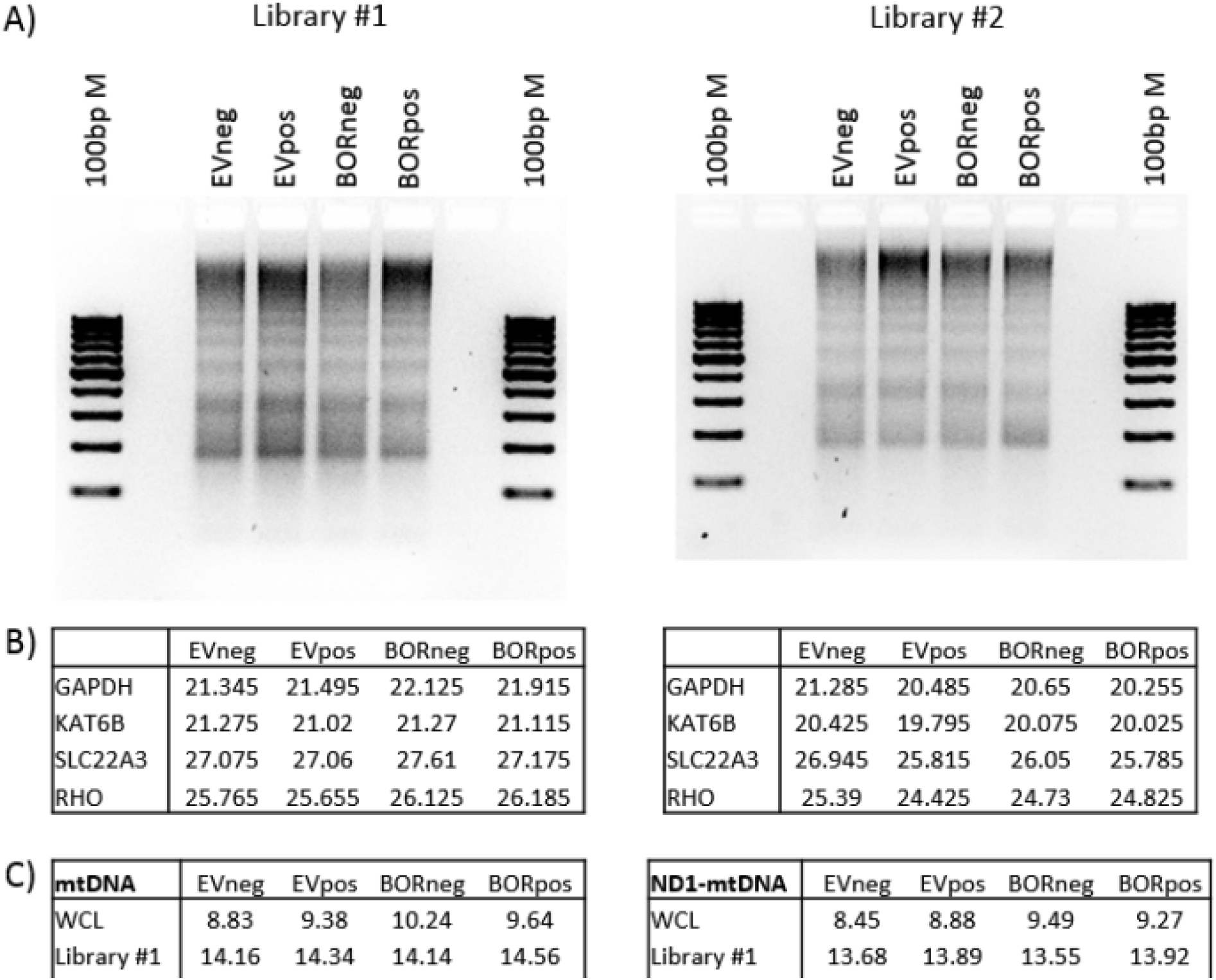
ATAC-seq library quality control. (A) Part of the DNA from each ATAC-seq library was analyzed by agarose gel to verify the presence of a nucleosome pattern. (B) and (C) Quantitative cycle (Cq) value for each target as assessed by qPCR on DNA from ATAC-seq libraries. B) Primers for GAPDH and KAT6B were used to amplify regions of open chromatin and SLC22A3 and RHO for regions of closed chromatin. (C) Two sets of primers (mtDNA and ND1-mtDNA) were used to amplify mtDNA. WCL: library prepared from whole cell lysate instead of isolated nuclei.

**Supplement 10.**
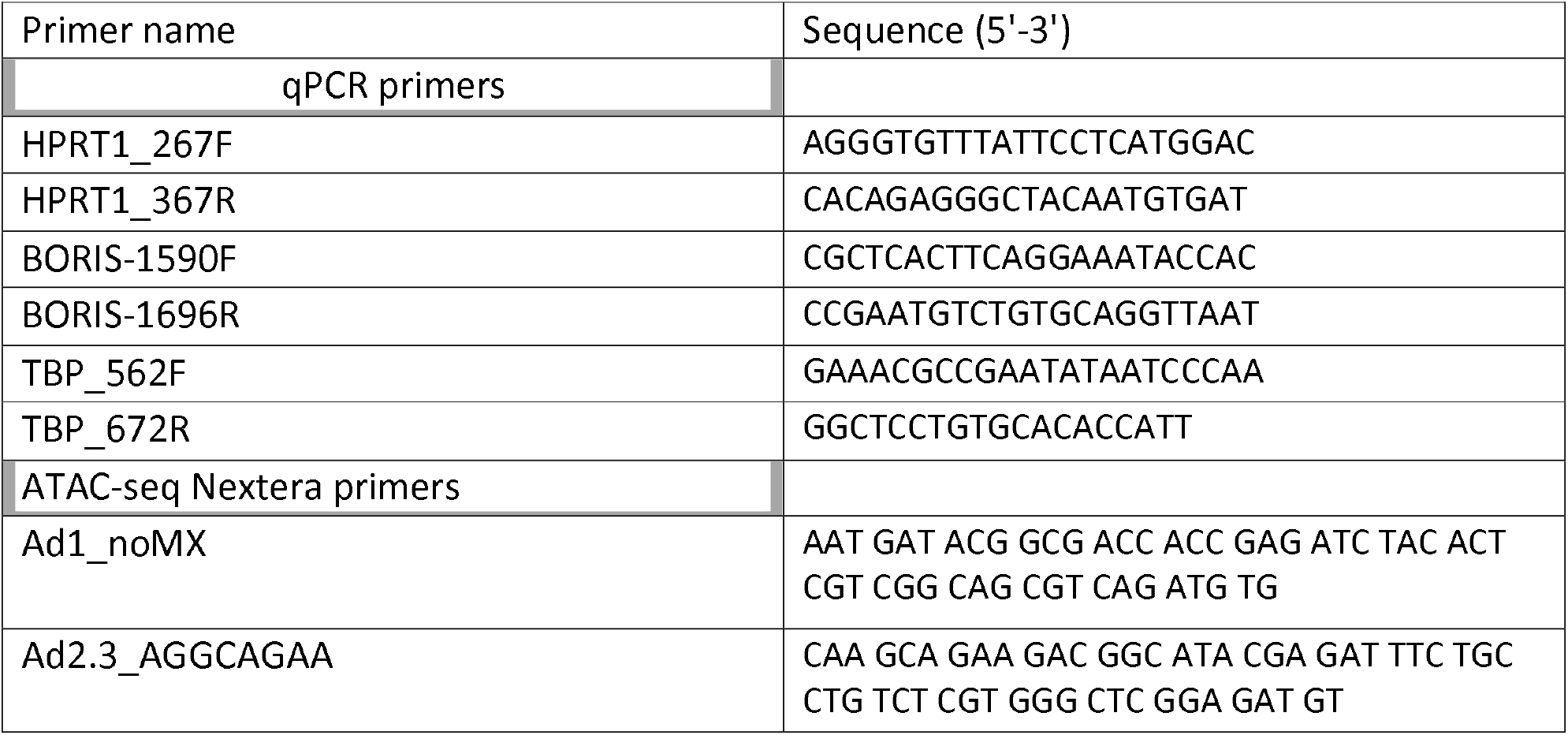
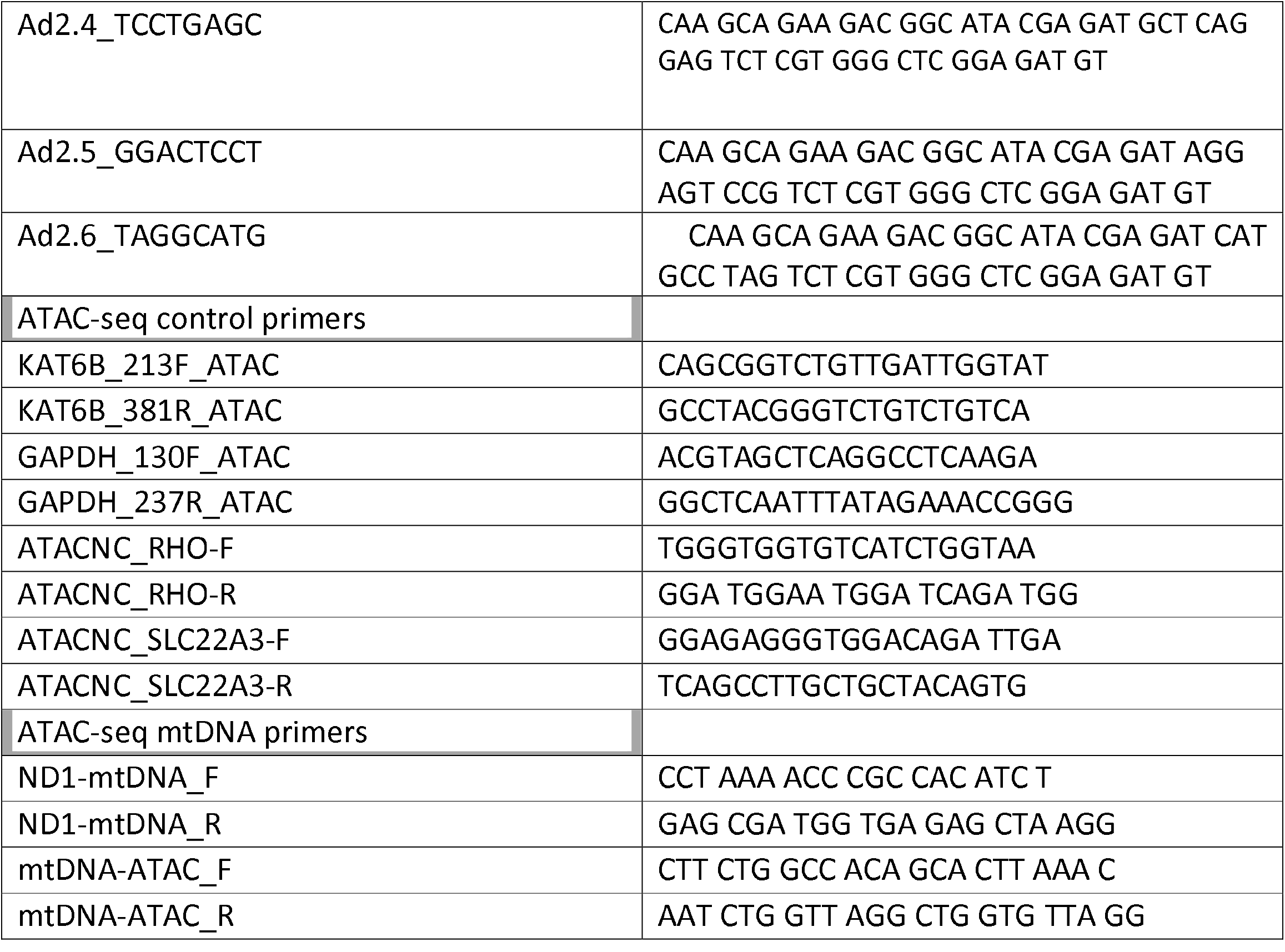
Primers table.

